# Developmental Genetics of Color Pattern Establishment in Cats

**DOI:** 10.1101/2020.11.16.385609

**Authors:** Christopher B. Kaelin, Kelly A. McGowan, Gregory S. Barsh

## Abstract

Intricate color patterns are a defining aspect of morphological diversity in the Felidae. We applied morphological and single-cell gene expression analysis to fetal skin of domestic cats to identify when, where, and how, during fetal development, felid color patterns are established. Early in development, we identify stripe-like alterations in epidermal thickness preceded by a gene expression pre-pattern. The secreted Wnt inhibitor encoded by Dickkopf 4 (*Dkk4*) plays a central role in this process, and is mutated in cats with the Ticked pattern type. Our results bring molecular understanding to how the leopard got its spots, suggest that similar mechanisms underlie periodic color pattern and periodic hair follicle spacing, and identity new targets for diverse pattern variation in other mammals.

## Main

Understanding the basis of animal color pattern is a question of longstanding interest for developmental and evolutionary biology. In mammals, markings such as cheetah spots and tiger stripes helped motivate theoretical models, such as the Turing reaction-diffusion mechanism, that have the potential to explain how periodic and stable differences in gene expression and form might arise from a uniform field of identical cells^1–4^. Reaction-diffusion and other mechanisms to account for periodic morphological structures have been implicated in diverse developmental processes in laboratory animals^5–12^, but much less is known about mammalian color patterns, largely because the most prominent examples occur in natural populations of wild equids and felids that are not suitable for genetic or experimental investigation.

In fish, color patterns involve direct interactions between pigment cells that are often dynamic, allowing additional pattern elements to appear during growth or regeneration^13–15^. By contrast, in mammalian skin and hair, melanocytes are uniformly distributed during development, and the amount and type of melanin produced is controlled later by paracrine signaling molecules within individual hair follicles^16–18^. Additionally, pattern element identity of an individual hair follicle, e.g. as giving rise to light- or dark-colored hair, is maintained throughout hair cycling and cell division, so that individual spots or stripes of hair apparent at birth enlarge proportionally during postnatal growth. Thus, periodic mammalian color patterns may be conceptualized as arising from a three-stage process: (1) Establishment of pattern element identity during fetal development; (2) Implementation of pattern morphology by paracrine signaling molecules produced within individual hair follicles; and (3) Maintenance of pattern element identity during hair cycling and organismal growth^17,19^.

Domestic cats are a useful model to study color patterns due to their accessibility, the genetic and genomic infrastructure, the opportunity for genomic and histological studies of tissue samples, and the diversity of pattern types^19–21^. The archetypal tabby pattern—regularly spaced dark markings on an otherwise light background—varies considerably in both form and color, and many of those varieties are similar to some wild felid species. In previous studies of domestic cats, we showed that *Endothelin 3* is expressed at the base of hair follicles in tabby markings, causes a darkening of the overlying hairs by increasing the production of black-brown eumelanin relative to red-yellow pheomelanin, and therefore plays a key role in implementation of tabby pattern^17^. However, tabby markings are apparent in developing hair follicles^17^, indicating that establishment of color pattern must occur at or before hair follicle development.

Here, we apply single cell gene expression analysis to fetal cat skin to investigate the developmental, molecular, and genomic basis of pattern establishment. We uncover a new aspect of epidermal development that signifies the establishment of pattern element identity and characterize signaling molecules and pathways associated with pattern establishment. We show that one of those signaling molecules encoded by *Dickkopf 4* (*Dkk4*) underlies a naturally occurring mutation that affects tabby pattern and plays a key role in the patterning process. Our work provides fundamental insight into the mechanisms of color pattern establishment as well as a platform for exploring the biology of periodic patterns more broadly.

## Results

### Patterns of epidermal morphology in developing cat skin

Trap-neuter-release programs have become a popular means of controlling feral cat overpopulation^22^. During breeding season, approximately half of all female feral cats are pregnant, and tissue from non-viable embryos can be recovered during spaying without compromising animal health or interfering with efforts to control feral cat overpopulation. We collected more than two hundred prenatal litters from feral cat, spay-neuter clinics across a range of developmental stages classified according to previous work on cats^23^ and laboratory mice^24^ (Supplementary Table 1).

Histochemical and morphometric analysis revealed a previously unknown aspect of epidermal development. At stage 13 (analogous to mouse embryonic day 11, E11), fetal skin is comprised of a uniform monolayer of epithelial cells that covers a pauci-cellular dermis (Fig. 1a). Approximately 16 days later at stage 16 (analogous to mouse E15), before epidermal differentiation and hair follicle morphogenesis, we noticed that the epidermis is organized into alternating regions that are either “thick” or “thin” (Fig. 1a, stage 16). Characterization of keratin expression and cell proliferation indicate that the thick and thin regions are fundamentally different from epidermal stratification that normally occurs later in development (Fig. 1b, 1c, Supplementary Note 1, Supplementary Table 1).

**Fig 1:**
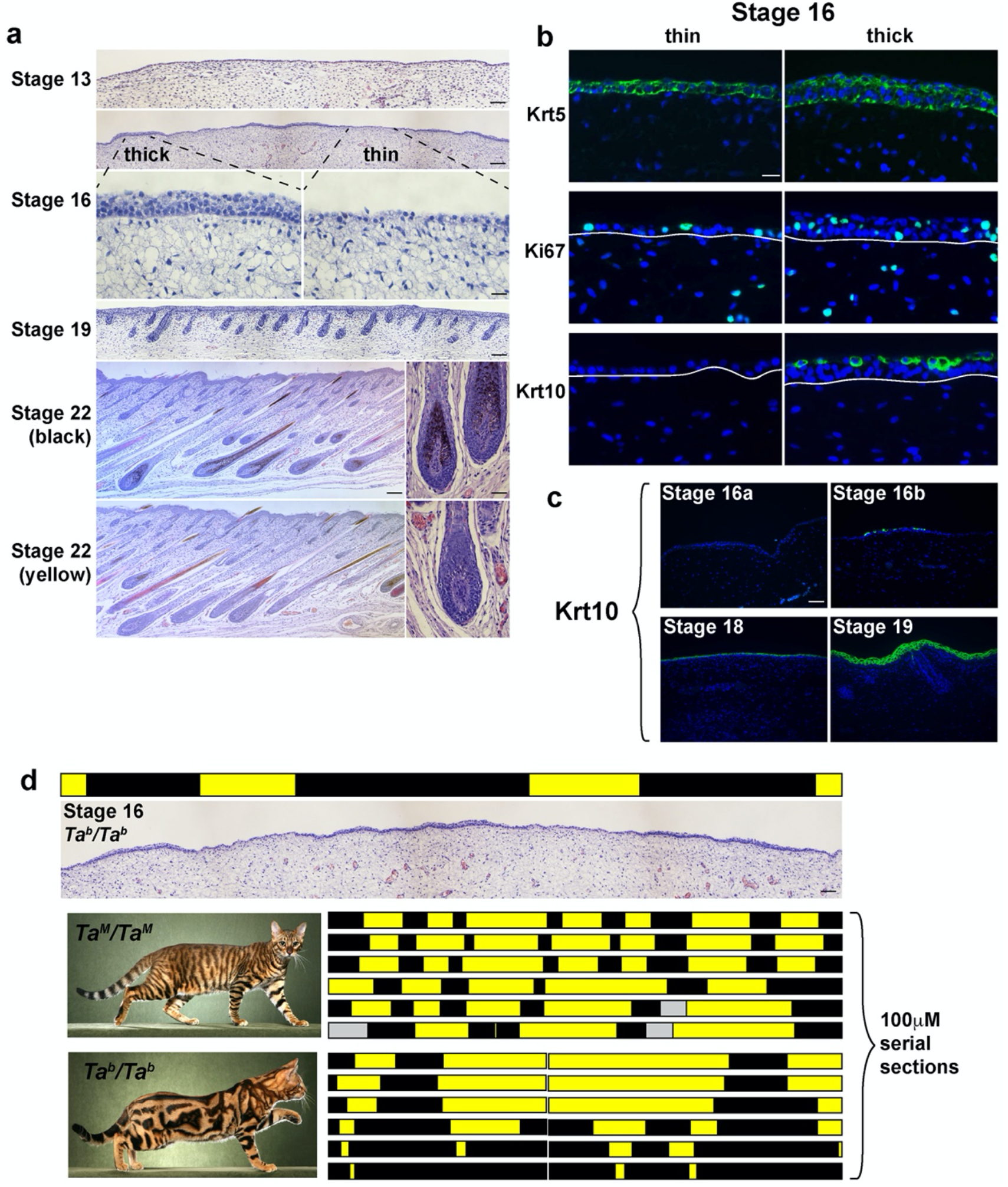
Patterns of epidermal thickening in fetal cat skin. **a**, Cat skin histology at different developmental stages (stage 16, bottom panels – high power fields of thick and thin epidermis; stage 22, right panels – high power fields of black and yellow follicles). **b**, Immunofluorescence for Krt5, Ki67 and Krt10 (green) in stage 16 skin sections (DAPI, blue; white lines mark dermal-epidermal junction). **c**, Krt10 immunofluorescence (green) on cat skin at different developmental stages (DAPI, blue). **d**, Topological maps based on skin histology (thin, yellow; thick, black) from *Ta*^*M*^/*Ta*^*M*^ and *Ta*^*b*^/*Ta*^*b*^ stage 16 embryos mimic pigmentation patterns observed in adult animals (left panels; gray bar - no data). Scale bars: a, stage 13 50 μM; stage 16 (top), Stage 19, stage 22 (left) 100 μM; stage 16 (bottom), stage 22 (right) 25 μM; b, 25 μM; c, d 50 μM.

By stage 22 (analogous to postnatal day 4-6 in laboratory mice), well-developed hair follicles are present that can be categorized according to the type of melanin produced (Fig. 1a), and that give rise to tabby pattern: dark markings contain mostly eumelanin, while the light areas contain mostly pheomelanin.

To investigate if epidermal thickening at stage 16 might be related to tabby patterns that appear later, we made use of natural genetic variation in *Transmembrane aminopeptidase Q*, (*Taqpep*). We showed previously that loss-of-function mutations in *Taqpep* cause the ancestral pattern of dark narrow stripes (*Ta*^*M*^/−, Mackerel) to expand into less well-organized large whorls (*Ta*^*b*^/*Ta*^*b*^, Blotched) (Fig. 1d)^17^. *Ta*^*b*^ alleles are common in most feral cat populations^25^, and we asked if the topology of stage 16 epidermal thickening is influenced by *Taqpep* genotype. Two-dimensional maps assembled from 100 μM serial sections of embryos of different genotype reveal that thick epidermal regions from *Ta*^*M*^/*Ta*^*M*^ embryos are organized into vertically oriented columns (black bars, Fig. 1d) separated by larger thin epidermal regions (yellow bars, Fig. 1d). By contrast, in *Ta*^*b*^/*Ta*^*b*^ embryos, the thick epidermal regions in the flank are broadened. A similar observation applies to epidermal topology and tabby patterns in the dorsal neck (Fig. 1d, Supplementary Fig. 1, Supplementary Note 1). Thus, embryonic epidermal topologies resemble tabby patterns in adult animals of the corresponding genotype, providing a morphologic signature of color pattern establishment before melanocytes enter the epidermis and before development of hair follicles. As described below, an associated molecular signature further refines the process to a 4-day window that spans stages 15 and 16, and delineates developmental substages (Supplementary Table 1) that track color pattern establishment more precisely.

### Single cell gene expression analysis of developing fetal skin

We hypothesized that the alternating thick and thin regions in fetal epidermis might arise from an earlier molecular pre-pattern. To explore this idea, we dissociated fetal cat skin, enriched for basal keratinocytes, and generated single-cell 3’ RNAseq (scRNAseq, 10x Genomics) libraries.

Among libraries that we sequenced and analyzed, 3 were from developmental stages (15a, 15b, 16a, Supplementary Table 1) occurring prior to or during the appearance of epidermal thickening. Uniform Manifold Approximation and Projection (UMAP) dimensionality reduction^26^ delineates different fetal skin cell populations as distinct cell clusters (Fig. 2a, Supplementary Table 3), but basal keratinocytes, identified by *Krt5* expression^27^, are the predominant cell type at all three stages (Supplementary Table 4, Supplementary Note 2). At stage 16a, unsupervised *k*-means clustering (*k*=9) is able to distinguish two subpopulations of basal keratinocytes based on differential expression of 604 genes (FDR<0.05), including the secreted inhibitors of Wnt signaling encoded by *Dickkopf 4* (*Dkk4,* p=6.50e-46, negative binomial exact test) and *Wingless Inhibitory Factor 1* (*Wif1*, p=5.81e-50, negative binomial exact test) (Supplementary Table 5). *Dkk4* and *Wif1* exhibit a similar extent of differential expression, 18-fold and 28-fold, respectively, but in the cells in which those genes are upregulated, the absolute levels of *Dkk4* are much higher than that of *Wif1*, 36 transcripts per cell compared to 2.2 transcripts per cell (Supplementary Table 5).

**Fig. 2:**
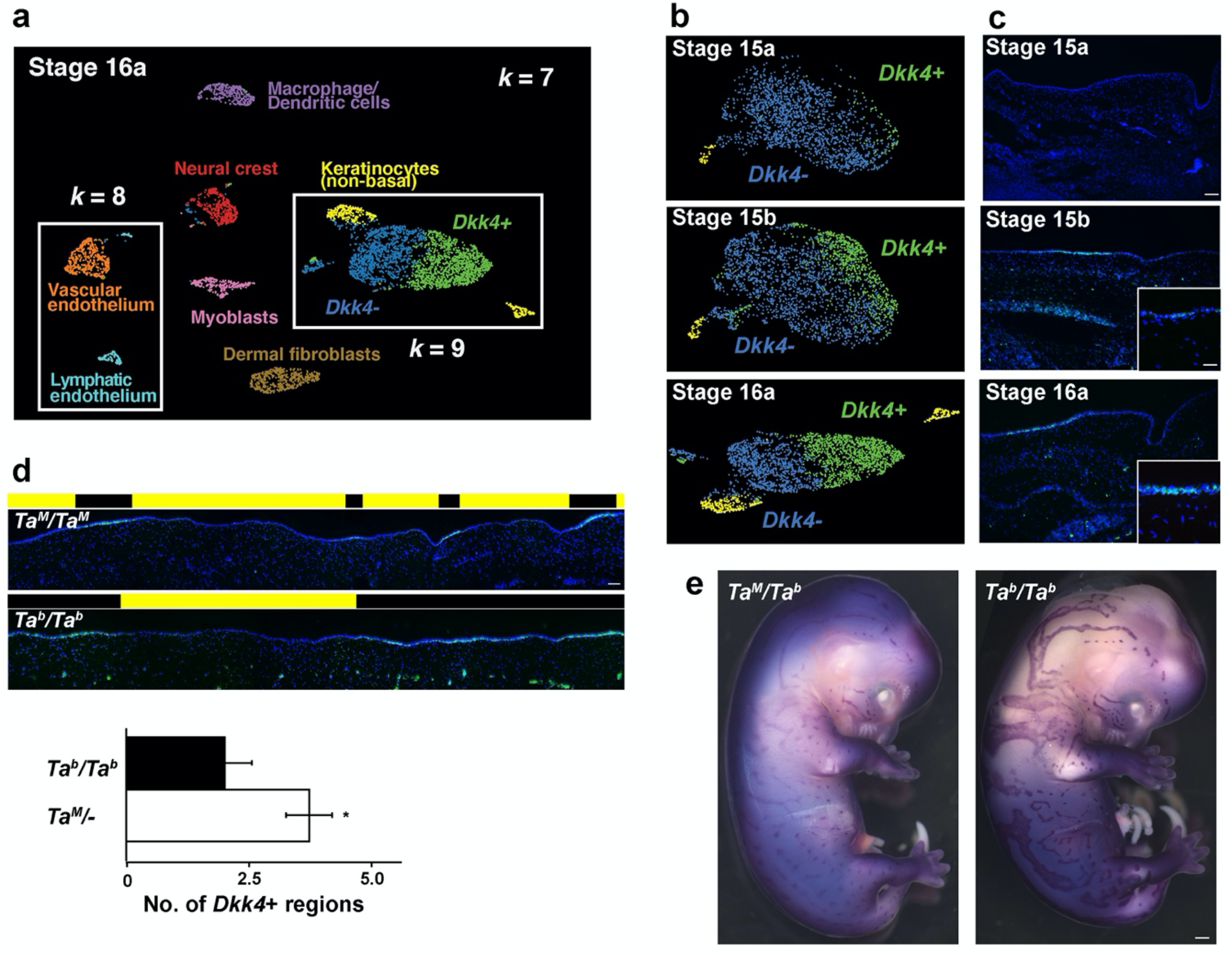
Pattern of gene expression in fetal cat skin determined by single-cell RNA sequencing and *in situ* hybridization. **a**, UMAP visualization of stage 16a cell populations colored by *k*-means clustering. Keratinocytes cluster as non-basal (yellow) and basal populations at *k*=7, with the latter split into subpopulations distinguished by high (green) and low (blue) *Dkk4* expression at *k*=9. Subpopulations of endothelium appear at *k*=8 (orange, cyan). **b**, Supervised (top and middle panels; stage 15a and 15b) and *k*-means (bottom panel, stage 16a) clustering at sequential developmental stages delineate a non-basal (yellow) and two basal keratinocyte populations, *Dkk4*-positive (green) and *Dkk4*-negative (blue). **c**, *Dkk4* expression (green) in fetal cat skin at corresponding stages (DAPI, blue; high power image, inset). **d**, *Dkk4* expression (green) in sections of stage 16 *Ta*^*M*^/ *Ta*^*M*^ and *Ta*^*b*^/*Ta*^*b*^ embryonic cat skin (DAPI, blue). Black and yellow blocks mark thick and thin regions, respectively. Number of *Dkk4*-positive regions (mean+/−sem) in 3.6mm of Stage 16 *Ta*^*M*^/− and *Ta*^*b*^/*Ta*^*b*^ embryonic cat skin from sections (n = 3-4 regions from at least two animals of each genotype; two-tail t-test). **e**, *Dkk4* expression (purple) in stage 16 *Ta*^*M*^/*Ta*^*b*^ and *Ta*^*b*^/*Ta*^*b*^ cat embryos. Scale bars: c, 50 μM; c inset, 25 μM; d, 50 μM; e, 1 mm.

Unsupervised clustering did not resolve different populations of basal keratinocytes at earlier stages, but a supervised approach based on differential expression of *Dkk4* (upper and middle panels, Fig. 2b, Supplementary Note 2) indicates that the same population apparent at stage 16a can also be recognized at stage 15a and 15b. As with stage 16a, the absolute levels of *Dkk4* expression are the highest of the differentially expressed genes, 22.3 and 99.2 transcripts per cell at stages 15a and 15b, respectively (Fig. 3c, Supplementary Table 5).

**Fig. 3.**
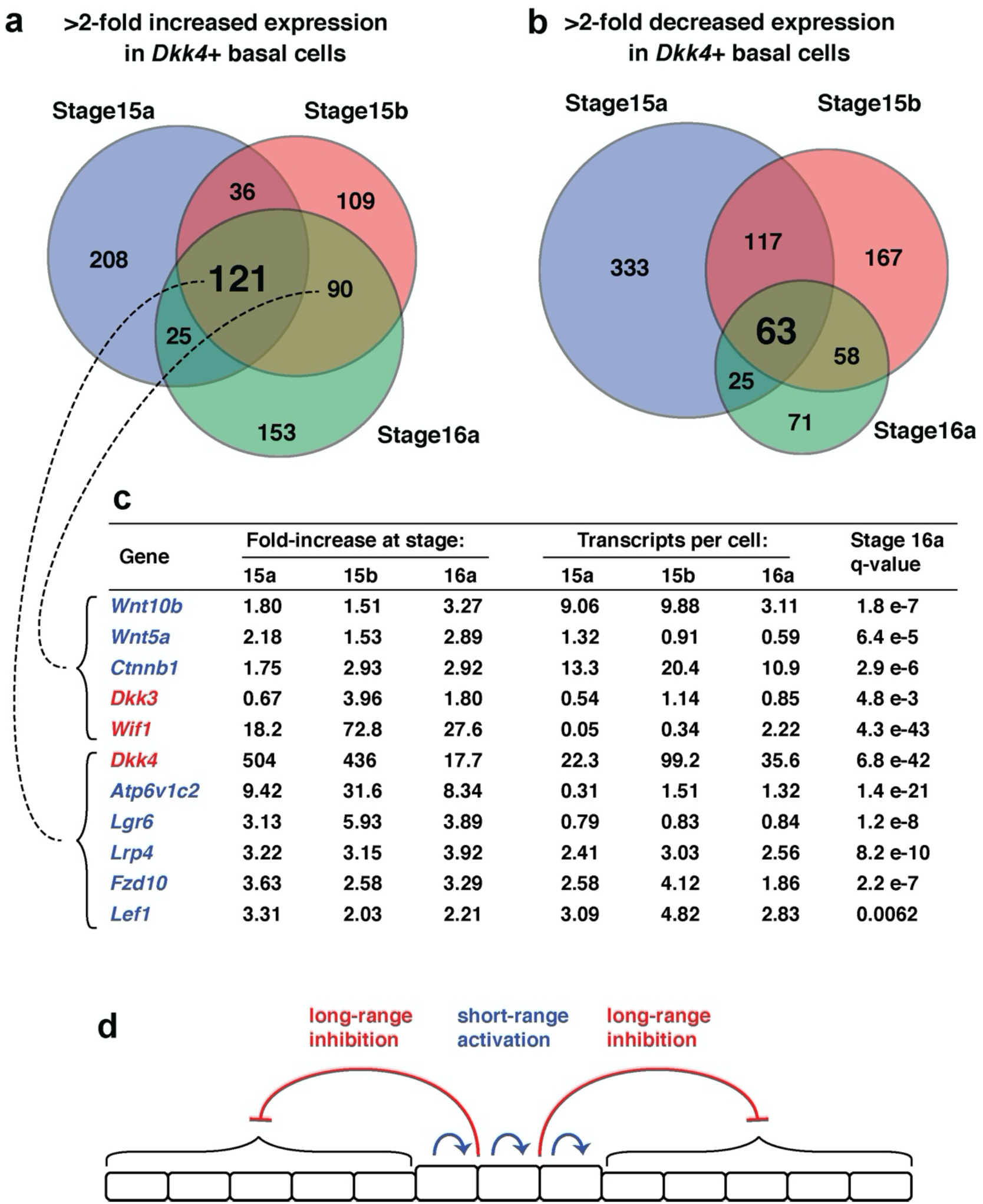
Single-cell RNA sequencing at successive stages of fetal skin development implicates WNT signaling-based reaction-diffusion in color pattern establishment. Overlap of gene expression profiles for genes with >2-fold increased (**a**) or decreased (**b**) expression in *Dkk4*-positive basal keratinocytes at stages 15a (blue), 15b (red), and 16a (green). (**c**) Gene expression metrics for WNT signaling genes with elevated expression in *Dkk4*-positive basal keratinocytes; q-values indicate the significance of differential expression between basal keratinocyte subpopulations at stage 16a (negative binomial exact test). Genes are colored according to their predicted role in either short-range activation (blue) or long-range inhibition (red) of WNT signaling in thick and thin epidermis, respectively, as presented in a model for color pattern establishment (**d**).

These observations on *Dkk4* in the scRNAseq data were confirmed and extended by *in situ* hybridization to fetal skin sections, and reveal that *Dkk4* is expressed in alternating subsets of basal keratinocytes at stage 15b, eventually marking the thick epidermal regions that are histologically apparent by stage 16a (Fig. 2c). Between stage 16a and 17, average size of the *Dkk4* expression domain becomes gradually smaller until it is only apparent in epidermal cells comprising the developing hair germ (Supplementary Fig. 2a, 2b, Supplementary Note 1).

As with the topology of epidermal thickening, we assessed the effect of *Taqpep* genotype on the pattern of *Dkk4* expression at stage 16, and observed a similar outcome: regions of *Dkk4* expression are fewer and broadened in *Ta*^*b*^/*Ta*^*b*^ embryos compared to *Ta*^*M*^/*Ta*^*M*^ embryos (Fig. 2d). Qualitatively, the effect is strikingly apparent from whole mount *in situ* hybridization (Fig. 2e, Supplementary Fig. 2c). These embryonic expression differences foreshadow the difference in adult color patterns between *Ta*^*M*^/− and *Ta*^*b*^/*Ta*^*b*^ animals. Thus, expression of *Dkk4* represents a dynamic molecular pre-pattern in developing skin that precedes and is coupled to the topology of epidermal thickening.

Examining the changes in basal keratinocyte gene expression more closely reveals an underlying transcriptional network that involves Wnt signaling (Fig. 3c, Supplementary Table 5). There are 927, 761, and 507 genes whose expression level changes >2-fold at stages 15a, 15b, and 16a, respectively, of which 121 upregulated and 63 downregulated genes are in common across all stages (Fig. 3a, 3b). Although the Wnt inhibitor encoded by *Dkk4* is the most highly expressed gene as noted earlier, other upregulated genes include *Atp6v1c2*, *Lgr6, Lrp4*, *Fzd10*, *and Lef1*, which represent a signature for increased responsiveness to Wnt signaling (Fig. 3c)^28,29^. At stages 15b and 16a, additional upregulated genes include *Wnt10b*, *Wnt5a*, *Ctnnb1*, and *Dkk3*. Overall, these patterns of gene expression suggest a reaction-diffusion model for establishment of color pattern in which *Dkk4*, *Dkk3*, and *Wif1* serve as long-range inhibitors, and *Wnt10b* and *Wnt5a* serve as short-range activators (Fig. 3d).

In laboratory mice, a Wnt—Dkk reaction diffusion system has been suggested previously to underlie periodic hair follicle spacing, based on patterns of gene expression and gain-of-function transgenic experiments^12,30^. In fetal cat skin, epidermal expression domains of *Dkk4* are much broader and appear earlier than those associated with hair follicle placode spacing (Supplementary Fig. 2a, 2b), although by stage 17, expression of *Dkk4* in cats is similar to expression of *Dkk4* in mice.

We compared genes that were differentially expressed in basal keratinocyte subpopulations during color pattern establishment in cats with those that are differentially expressed between hair follicle placodes and interfollicular epidermis in mice, based on a previously established 262 gene signature for primary hair follicle placodes^28^. Of the 121 genes that we identified as a signature for color pattern establishment in *Dkk4*-positive basal keratinocytes, 26 (21.5%) were shared with the primary hair follicle placode signature (Supplementary Table 5). Thus, the gene expression pre-pattern for color pattern establishment occurs prior to and foreshadows an overlapping pattern in hair follicle placodes.

### A *Dkk4* mutation in domestic cats

The mackerel stripe pattern represents the ancestral state; in addition to the blotched pattern caused by *Taqpep* mutations (Fig. 1d), several additional pattern types are recognized in domestic cats for which the genetic basis is uncertain. For example, periodic dark spots as seen in the Egyptian Mau or Ocicat breeds (Fig. 4a) are only observed in *Ta*^*M*^/− animals^19,31^, but the genetic basis of Mackerel vs. Spotting is not known. Another locus, *Ticked*, named for its ability to prevent dark tabby markings and thereby showcase hair banding patterns across the entire body surface, has been selected for in breeds with a uniform appearance such as the Abyssinian, Burmese, or Singapura (Fig. 4a)^32^. However, *Ticked* is also thought to be responsible for the so-called “servaline” pattern of spotted Savannah cats (Fig. 4d), in which large dark spots are reduced in size and increased in number. *Ticked* was originally thought to be part of an allelic series that included *Ta*^*M*^ and *Ta*^*b*^ ^33^, but was subsequently mapped to an independent locus on chrB1, and recognized as a semidominant derivative allele, *Ti^A^*, obscuring tabby markings except on the legs and the tail when heterozygous, and eliminating tabby markings when homozygous^19,34^.

**Fig. 4:**
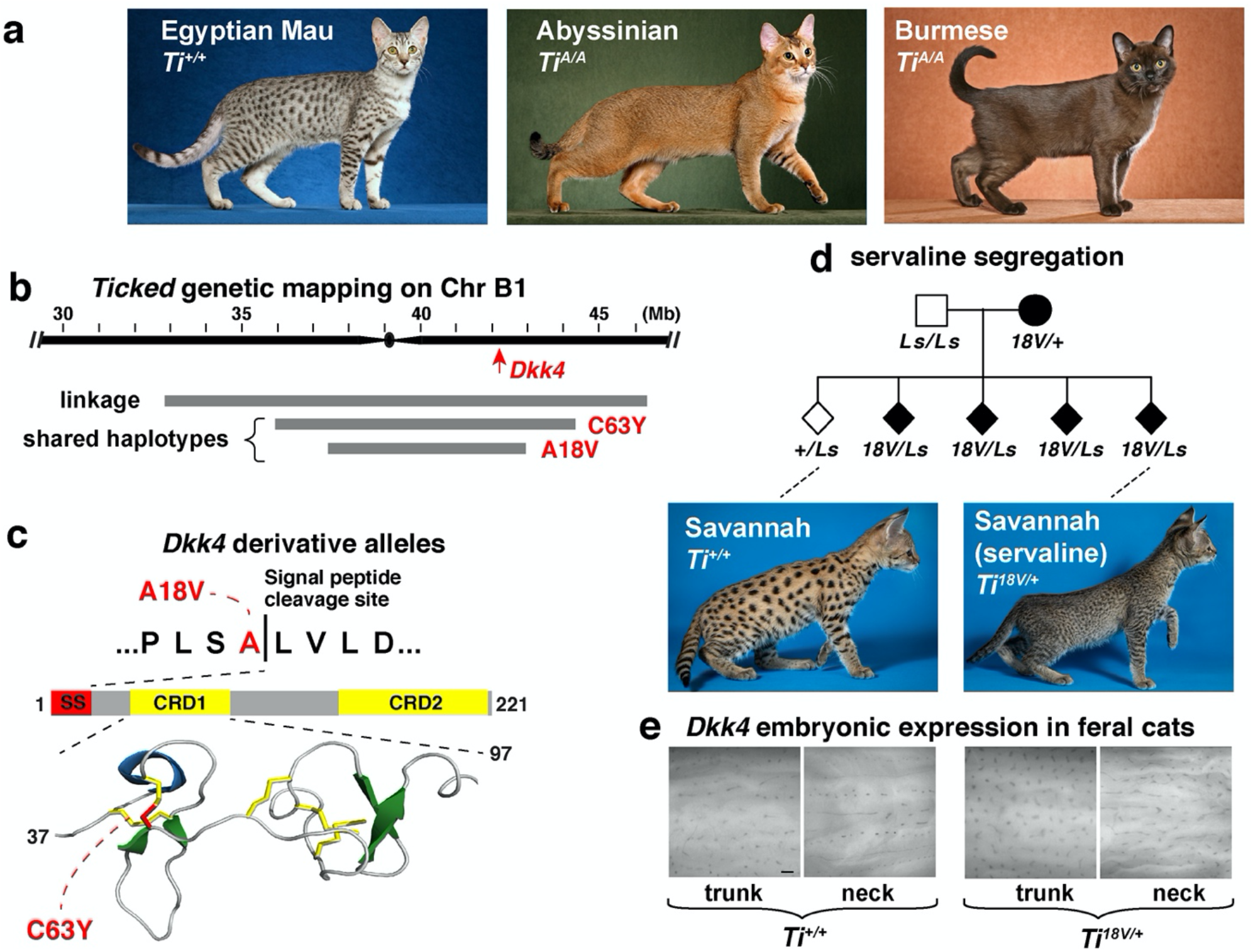
*Dkk4* mutations alter *Dkk4* expression patterns in fetal skin and coat color patterns in adult cats. **a**, Cat breeds selected for Spotted (Egyptian Mau) or for Ticked (Abyssinian and Burmese) color patterns. **b**, *Ticked* genetic interval delineated by linkage^19^ or shared haplotypes among cats with *Dkk4* mutations. Coordinates are based on the felCat9 assembly. **c**, *Dkk4* coding mutations (red) at the signal peptide cleavage site (p.Ala18Val) or a cysteine residue (p.Cys63Tyr) involved in disulfide bridge and cysteine knot formation within cysteine rich domain 1 (CRD1). **d**, Cosegregation of *Dkk4* p.Ala18V and the servaline color pattern in Savannah cats. **e**, Altered *Dkk4* expression (dark markings) in whole mount preparations from stage 17 fetal skin. Scale bar in e, 300 μM.

*Ticked* maps to a region of low recombination close to the centromere, and has therefore been difficult to identify by linkage analysis. Furthermore, we observed that some cats thought to be homozygous for *Ticked*, including Cinnamon, the source of the feline reference sequence, carried two haplotypes across the linkage region, suggesting the existence of multiple *Ti*^*A*^ mutations. We hypothesized that one or more of the genes that was differentially expressed between basal keratinocyte subpopulations might represent *Ticked*, and asked whether any differentially expressed gene satisfied two additional criteria: (1) a genetic map position close to or coincident with *Ticked*; and (2) carrying a deleterious variant found at high frequency in breeds with a Ticked phenotype. Of the 602 differentially expressed genes at stage 16a, 4 are located in a relatively small domain, ~230 kb, that overlaps with the *Ticked* linkage interval (Fig. 4b). One of these genes, *Dkk4*, exhibits a pattern of variation and association consistent with genetic identity as *Ticked*.

We surveyed the 99 Lives collection of domestic cat whole genome sequence data^35^, and identified two nonsynonymous variants in *Dkk4*, p.Ala18Val (felCat9 chrB1:42620835c>t) and p.Cys63Tyr (felCat9 chrB1:42621481g>a), at highly conserved positions (Supplementary Fig. 3) that were present only in breeds with obscured tabby markings (Abyssinian, n=4; Burmese, n=5; Siamese n=1). The p.Ala18Val variant is predicted to impair signal peptide cleavage, which normally occurs between residues 18 and 19, and the p.Cys63Tyr variant is predicted to disrupt disulfide bonding in a cysteine knot structure referred to as CRD1^36^ (Fig. 4c, Supplementary Fig. 3). These are the only coding alterations predicted to be deleterious in any of the 4 genes that are differentially expressed in basal keratinocyte subpopulations (Supplementary Table 6), and are therefore strong candidates for *Ticked*.

Association analysis of additional DNA samples across breeds (n=115) and within breeds (n=238) (Table 1) provides further support that variation in *Dkk4* causes *Ticked* (Table 1, Supplementary Table 7, Supplementary Note 3). All breed cats in which *Ticked* is required (Abyssinian, Singapura) carried the p.Ala18Val or p.Cys63Tyr *Dkk4* variants, the vast majority as homozygotes or compound heterozygotes. By contrast, in breeds in which tabby markings are required (Egyptian Mau, Ocicat, Bengal), none carried a derivative *Dkk4* allele. Lastly, in some breeds (Oriental Longhair, Oriental Shorthair) and in non-breed cats, either Ticked or non-Ticked phenotypes are observed, and are perfectly correlated with the presence or absence of the p.Ala18Val *Dkk4* variant.

**Table 1.**
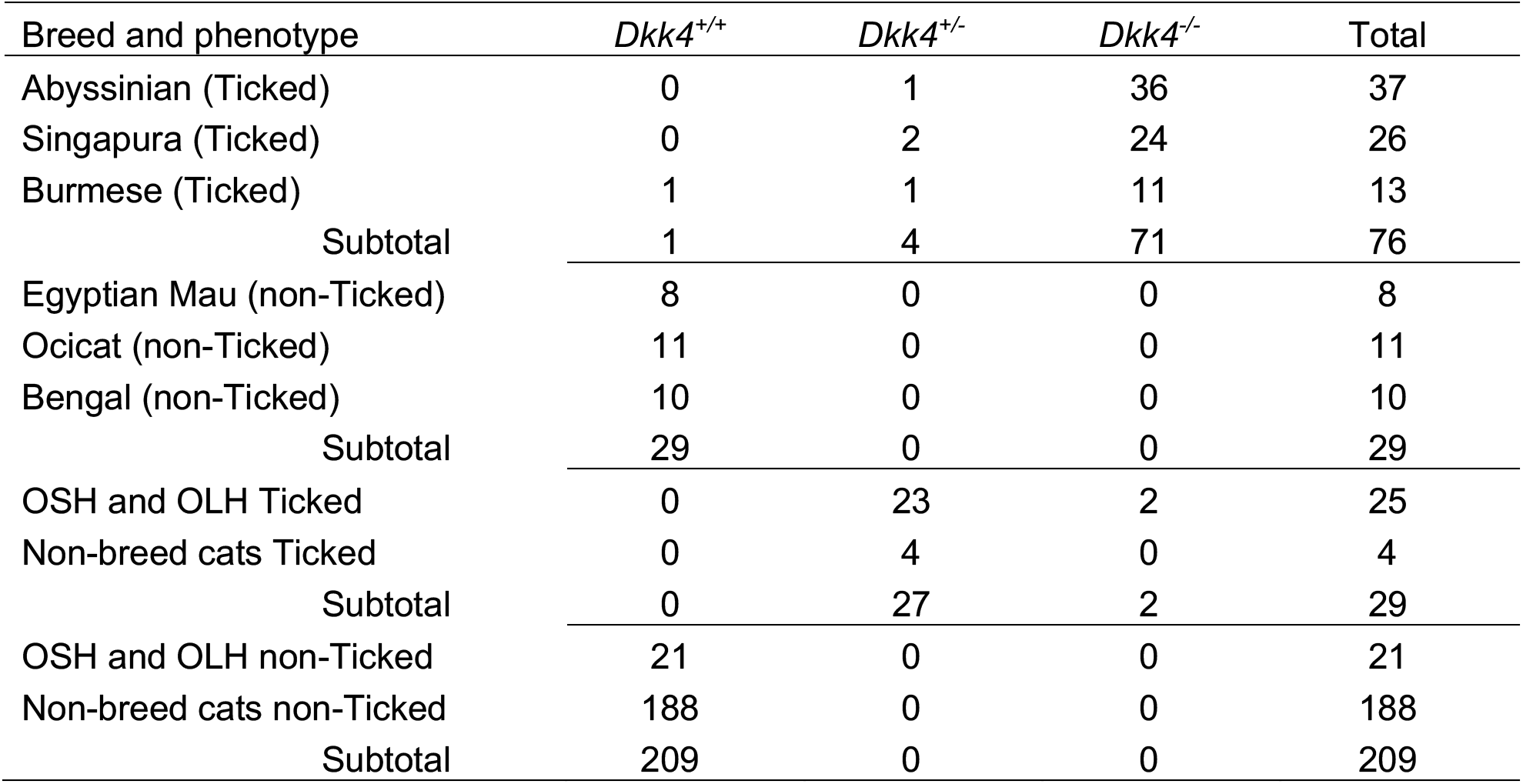
*Dkk4* genotypes in Ticked and non-Ticked cats.

Haplotype analysis indicates that the p.Ala18Val and p.Cys63Tyr variants occurred on common and rare haplotypes, respectively (Supplementary Fig. 4a). We identified several small pedigrees where one or both of the *Dkk4* variants are present (Fig. 4d, Supplementary Fig. 4b, 4c), and observed segregation patterns consistent with genetic identity of *Dkk4* and *Ticked*. A pedigree of Savannah cats in which the servaline phenotype cosegregates with the p.Ala18Val variant is of particular interest (Fig. 4d): rather than an apparent suppression of tabby markings as in the Abyssinian and Burmese breeds, *Ticked* alters the number and size of tabby markings.

Unlike *Taqpep*, variation for *Ticked* in California feral cat populations is rare (Extended Data Table 4); however, one of our fetal skin samples was heterozygous for the p.Ala18Val variant, enabling us to examine the effect of *Dkk4* on its own expression. As depicted in Fig. 4e, whole mount *in situ* hybridization of a *Dkk4* probe to fetal skin of *Ti*^*+/+*^ and *Ti*^*+/A18V*^ individuals yields patterns that are remarkably similar to the adult Savannah and servaline color markings, respectively. Taken together, these results confirm that the effect of *Ticked* is not to mask dark tabby markings (as it appears to do in the Abyssinian and Burmese breeds) but to affect pattern establishment such that the regions that express *Dkk4* in fetal keratinocytes, and the eventual adult markings, become smaller and more numerous.

## Discussion

Key elements of reaction-diffusion models as applied to biological pattern are the existence of diffusible signaling molecules that interact with one another to achieve short-range activation and long-range inhibition^37^. Our work identifies molecular candidates for this process in establishment of tabby pattern, suggests that similar mechanisms underlie periodic color patterning and periodic hair follicle spacing, and provides a genomic framework to explore natural selection for diverse pattern types in wild felids.

Development of tabby pattern is different, and arguably simpler, than developmental systems that involve extensive cell movement or cytoplasmic protrusions, as in zebrafish^13,15,38^. Additionally, establishment of tabby pattern is completely and remarkably uncoupled from the implementation events that follow days and weeks later. In particular, epidermal expression of *Dkk4* is dynamic, marking areas of fetal skin that give rise to hair follicles that later produce dark pigment throughout successive hair cycles. This implies that *Dkk4*-expressing keratinocytes acquire time-sensitive epigenetic changes that are later incorporated into hair follicles, and that ultimately determine whether the underlying dermal papilla releases paracrine agents that darken or lighten the hair.

The molecular and the morphologic signatures of tabby pattern establishment recede by stage 17 as hair follicle placodes start to form. Nonetheless, the two processes share important features. A Wnt-Dkk reaction-diffusion system has been proposed to underlie hair follicle spacing^12^, and similarity between the transcriptional profiles described here and those in hair follicle placodes^28^ suggests the two processes could be mediated by similar mechanisms. Previous work on hair follicle spacing involved a gain-of-function perturbation in which overexpression of *Dkk2* in incipient hair follicles caused both a reduced number and a clustered distribution of follicles^12^. Our work on *Dkk4* mutations yields reciprocal results: the p.Ala18Val and p.Cys63Tyr alleles are loss-of-function and, in servaline cats, are associated with both an increased number and reduced size of dark spots.

In domestic cats, the servaline phenotype is one of several examples demonstrating that the effect of *Dkk4* depends on genetic background and genetic interactions, as exemplified by the different appearance of the Abyssinian and Burmese breeds, and by interaction of the *Ticked* (*Dkk4*) and *Tabby* (*Taqpep*) genes^19,39^. The effects of *Taqpep* on visualization of Blotched vs. Mackerel are apparent only in a non-Ticked background, i.e. *Dkk4* is epistatic to *Taqpep*, which encodes an aminopeptidase whose physiologic substrates have not been identified^17^. *Taqpep* is expressed in developing skin^17^, and our observations are consistent with a model in which Taqpep-dependent cleavage restricts the range and/or activity of Dkk4, representing a potential new mechanism for modifying Wnt signaling.

Variation in *Taqpep* is associated with pattern diversity in the cheetah as well as in the domestic cats^17^, and the same may be true for *Dkk4*. In fact the servaline pattern was first described in 1839 as characteristic of a new species, *Felis servalina*, later recognized as a rare morph of the Serval^40^. We examined *Dkk4* predicted amino acid sequences for 29 felid species including the Serval, and identified a number of derived variants (Supplementary Fig. 5). Although none of the derived variants represents an obvious loss-of-function, several are predicted to be moderately deleterious (Supplementary Table 9), and a wider survey of *Dkk4* variation in Servals could reveal additional mutations that explain the spotting pattern of *Felis servalina*.

More broadly, genetic components of the molecular signature we identified for color pattern establishment are potential substrates for natural selection and evolution of pattern diversity in wild felids and other mammals^41^. For example, complex patterns such as rosettes in the jaguar or complex spots in the ocelot may represent successive waves of a Wnt-Dkk system, similar to what has been proposed for hair follicle spacing^12^. Although the action of *Dkk4* in domestic cats appears limited to color pattern, subtle pleiotropic effects of *Dkk4* inactivation that reduce fitness may not be apparent. If so, regulatory variation in candidate genes identified here may be substrates for diversifying selection during felid evolution.

Our work exemplifies the potential of companion animal studies to reveal basic insight into developmental biology, and builds on a history in model organisms such as laboratory mice, in which coat color variation has been a fruitful platform for studying gene action and interaction. The Ticked, Tabby, and Blotched phenotypes represent a fraction of the pattern diversity that exists among domestic cat breeds and interspecific hybrids, which remain a continued resource to explore how spots and stripes form in nature.

## Methods

### Biological samples

Embryonic cat tissues were harvested from incidentally pregnant, feral cats at three different spay-neuter clinics in California. Tissues were processed within 18 hours of the spaying procedure and embryos were staged based on crown-rump length and anatomic development (Supplementary Table 1)^23^.

Genomic DNA for genetic association studies was collected with buccal swabs (Cyto-Pak, Medical Packaging Corporation) at cat shows or by mail submission from breeders, and extracted using a QIAamp DNA kit (Qiagen). Feral cat DNA was obtained from gonadal tissue or embryonic tail after spay/neuter surgeries, as described previously^17^. Genotyping is based on Sanger sequencing of PCR amplicons for *Dkk4* p.Ala18Val (GAGCTGAGAAGGTCAAGGTGA, GTGGGTACTTGTGCCATTCC), *Dkk4* p.Cys63Tyr (CCACTGTGATTTGGCTTCCT, CAGTCCCACAGGGGTTTATG), and *Taqpep* p.Trp841X (GCCTTCGGAAGTGATGAAGA, ACTTCAGATTCCGCCACAAC).

Sample collection and processing was conducted in accordance with a protocol approved by the Stanford Administrative Panel on Laboratory Animal Care.

### Histology and immunofluorescence

Embryonic cat tissues were fixed in 4% paraformaldehyde (Electron Microscopy Sciences). 5 μM paraffin embedded sections were mounted on Superfrost Plus glass slides (Fisher Scientific, Pittsburgh, PA) and stained with hematoxylin and eosin. Immunofluorescence for Krt5 (Biolegend), Krt10 (Abcam) and Ki67 (Abcam) was carried out on 5 μ M sections after antigen retrieval using 0.01 M citrate buffer, pH 6 in a pressure cooker. Krt5-stained sections were incubated with goat anti-rabbit Alexa488 antisera (Jackson ImmunoResearch Laboratories). Krt10 and Ki67 stained sections were incubated with goat anti-rabbit biotinylated antisera (Jackson ImmunoResearch Laboratories), Vectastain Elite Avidin-Biotin complex reagent (Vector Labs), and Tyramide-Cy3 amplification reagent (PerkinElmer). Sections were stained with ProLong antifade reagent with DAPI (Invitrogen). All photomicrographs are representative of at least three animals at each developmental time point. Precise genotypes are provided in the text and figures, except in instances where *Ta*^*M*^/*Ta*^*M*^ and *Ta*^*M*^/*Ta*^*b*^embryos were used together, which are designated as *Ta*^*M*^/−.

### *In situ* hybridization

A 257 bp digoxigenin-labeled RNA probe was generated from a region spanning exons 1-3 of cat *Dkk4* using a PCR-based template (cDNA from Stage 16 cat embryonic epidermal cells; CCCTGAGTGTTCTGGTTTT, AATATTGGGGTTGCATCTTCC) and *in vitro* transcription (Roche Diagnostics). 10 μ M paraffin-embedded sections were deparaffinized with xylenes and antigen retrieval using 0.01 M citrate buffer, pH 6 in a pressure cooker was carried out. Sections were treated with Proteinase K (Sigma), hybridized with 150 ng/ml riboprobe overnight at 60 degrees, and incubated with anti-digoxigenin antibody conjugated with horseradish peroxidase (Roche Diagnostics) and Tyramide-Cy3 amplification reagent (PerkinElmer). Sections were stained with ProLong antifade reagent with DAPI (Invitrogen).

Samples for whole mount *in situ* hybridization were treated with Proteinase K (Sigma), neutralized with 2 mg/ml glycine, fixed in 4% paraformaldehyde with 0.1% glutaraldehyde (Electron Microscopy Sciences), treated with 0.1% sodium borohydride (Sigma), and hybridized with 0.5 μg/ml *Dkk4* riboprobe at 60 degrees overnight. Embryos were subsequently treated with 2% blocking reagent (Roche Diagnostics) in maleic acid buffer with Tween-20 (MABT), incubated with an alkaline phosphatase-conjugated anti-digoxigenin antibody (Roche Diagnostics) overnight at 4 degrees, and developed 3-6 hours in BM Purple (Sigma). Nuclear fast red (Sigma) was used to stain 5 mM sections in Supplementary Fig. 2.

For analysis of *Dkk4* expression in fetal skin of *Ta*^*M*^/*Ta*^*M*^ and *Ta*^*b*^/*Ta*^*b*^ embryos (Fig. 2D), we quantified the difference between the two genotypes by counting the number of *Dkk4*-positive regions in 3.6 mm longitudinal segments of flank epidermis from sections of *Ta*^*M*^/*Ta*^*M*^ (n=4 epidermal segments from three embryos) and *Ta*^*b*^/*Ta*^*b*^ (n=3 epidermal segments from two embryos) embryos.

### Tissue preparation for single-cell RNA sequencing

A single embryo at each of three development stages (Stage 15a, 15b, and 16a) was used to prepare a dissociated cell homogenate for single-cell RNA sequencing. Embryos were treated with Dispase (Stem Cell Technologies) and the epidermis was removed with gentle scraping using a No.15-blade scalpel. Epidermal sheets were dissociated with trypsin (Gibco) and gentle agitation. Red blood cells were removed with a red blood cell lysis solution (Miltenyi Biotec). Single-cell suspensions were incubated with a rat anti-CD49f antibody conjugated with phycoerythrin (ThermoFisher Scientific, clone NKI-GoH3), followed by anti-phycoerythrin magnetic microbeads and enriched using a column-magnetic separation procedure (Miltenyi Biotec). 10^6^ fetal skin cells were processed for each single-cell RNA sequencing library.

### Single-cell RNAseq

3’ single-cell barcoding and library preparation were performed using 10x Genomics single-cell RNA sequencing platform (10x Genomics). Libraries were constructed with version 2 (stage 16a) or version 3 (stage15 a/b) chemistry and sequenced on a single HiSeqX (Illumina) lane to generate 400-450 million paired-end 150 bp reads (Supplementary Table 4).

Barcoded reads were aligned to the domestic cat genome assembly (felCat9, NCBI Felis catus Annotation Release 104) using the 10x Genomics CellRanger (v3.1) software. Annotations for 3’ untranslated regions were extended by 2 kb for all transcripts except when an extension overlapped a neighboring gene. The ‘cellranger count’ pipeline was used to output feature-barcode matrices, secondary analysis data, and browser visualization files. The ‘cellranger reanalyze’ function was used to exclude 247 cell cycle genes and *XIST* to minimize cell-cycle and sex associated variation in downstream analysis.

*k*-means clustering performed by the ‘cellranger count’ pipeline on the stage 16a data set was used to delineate cell populations (Supplementary Note 2), progressively increasing *k* until the basal keratinocyte population was split into distinct groups (at *k*=9). Differential expression analysis between cell populations defined by clustering was measured using Cell Ranger’s implementation of sSeq^42^, which employs a negative binomial exact test.

For stages 15a and 15b, the ‘cellranger aggr’ function was used to aggregate expression matrices from each stage. Aggregation resulted in co-clustering of stage15a (n=59) and 15b (n=774) *Dkk4*-positive cell populations in UMAP projections that failed to cluster at stage 15a when expression matrixes were analyzed independently. Differential expression analysis between high and low expressing *Dkk4* populations was measured using Cell Ranger’s implementation of sSeq, as described above.

Genomic coordinates for the 604 genes differentially expressed at stage 16a were determined using the felCat9 genome assembly feature table file (www.ncbi.nlm.nih.gov/assembly/GCF_000181335.3/). Functional annotation was performed using Panther (pantherdb.org) and DAVID (david.ncifcrf.gov) annotation tools.

### *Ticked* genetic analysis

Whole genome sequence data from 57 cats in the 99 Lives dataset were used for variant detection. Variant calling and annotation (NCBI Felis catus Annotation Release 103) were performed with Platypus^43^ and SNPeff^44^, respectively, after alignment to the domestic cat genome (felCat8) with BWA. SNPsift was used to identify protein coding variants in *Vdac3*, *Polb*, *Dkk4*, and *Plat*.

Beagle^45^ v4.1 was used to infer haplotypes across a 15 Mbp interval from chrB1:30,000,000-45,000,000 (felCat8). Phased SNPs were thinned to 1 site/5 kb with VCFtools and converted to felCat9 assembly coordinates with the UCSC LiftOver tool. The color coding for reference (blue) and alternate (yellow) alleles (Suppelmentary Fig. 4a) reflect the felCat8 assembly, but the genomic coordinates presented are converted to the felCat9 assembly.

Combined annotation dependent depletion^46^ (CADD, v1.4) was used to score the deleteriousness of coding variants within *Dkk4*, after converting to orthologous positions in the human genome (GRCh38/Hg38). SignalP-5.0 was used to predict signal peptide cleavage for the p.Ala18Val variant, and the PyMol browser (v2.3.3) was used to visualize and display the protein structure of the CRD1 domain of Dkk4 (Fig. 4c), using the N-terminal region solution structure of recombinant human Dkk4 protein^36^ (5O57, Protein Data Bank).

## Data availability

Raw and processed single-cell RNAseq datasets have been deposited in GEO (accession number: GSE152946). GEO files include unfiltered feature-barcode matrices in HDF5 format, output by the CellRanger pipeline, and Illumina fastq files for stage 15a, 15b, and 16a single-cell RNAseq.

## Acknowledgments

The authors thank Hermogenes Manuel for technical assistance, Anthony Hutcherson and other members of The International Cat Association and Cat Fancy Association for assistance with breed sample collection, Trap-Neuter-Release programs in California for assistance with feral sample collection, Valerie Smith, Adriana Kajon, and Gulnaz Sharifzyanova for providing material from OSH, Singapura, and Savannah pedigrees, respectively, Helmi Flick and Jamila Agaeva for domestic and Savannah cat photographs, respectively. This research has been supported in part by the HudsonAlpha Institute for Biotechnology, and by a grant from the National Institutes of Health to G.S.B (AR-067925).

## Authors contributions

C.B.K., K.A.M., and G.S.B. conceived the project, designed experiments, and wrote the paper. C.B.K., and K.A.M. performed experiments and analyzed data; G.S.B. procured funding.

## Supplementary materials

Supplementary Note 1: Embryonic staging and characterization of epidermal morphology.

Supplementary Note 2: Single-cell gene expression experiments and analyses.

Supplementary Note 3: Genetic evaluation of *Ticked*.

Supplementary Fig. 1. Topological maps of dorsal neck sin from Stage 16.

Supplementary Fig. 2. *Dkk4* expression in embryonic cat skin.

Supplementary Fig. 3. Evolutionary constraint and functional prediction for cat Dkk4 variants associated with *Ticked*.

Supplementary Fig. 4. *Dkk4* variants occur on extended haplotypes and co-segregate with *Ticked* in pedigrees.

Supplementary Fig. 5. Protein alignment of predicted Dkk4 protein sequence from 29 felid species

Supplementary Table 1. Stages of embryonic and fetal development in the domestic cat.

Supplementary Table 2. Histologic and molecular features of thick epidermal domains in cat as compared to developing hair follicle placodes and “normal” epidermal differentiation.

Supplementary Table 3. Gene markers used for identification of UMAP clusters

Supplementary Table 4. scRNAseq summary statistics

Supplementary Table 5. List of differentially expressed genes from scRNAseq and overlap with hair follicle placode (separate spreadsheet).

Supplementary Table 6. *Dkk4* coding variants detected in 57 cats.

Supplementary Table 7. *Dkk4* alleles categorized according to breed and phenotype.

Supplementary Table 8. Basal keratinocyte subpopulation cell number at different *Dkk4* expression thresholds.

Supplementary Table 9. *Dkk4* variants and predicted impact (CADD) in the Felidae.

