## Supplemental Material for "Developmental Genetics of Color Pattern Establishment in Cats"

**Supplementary Material for Kaelin, McGowan, and Barsh**

Supplementary Note 1: Embryonic staging and characterization of epidermal morphology

Previous work has classified the domestic cat's 63-day gestation into 22 developmental stages [1]. Supplementary Table 1 depicts the criteria we used for staging, including size (crown-rump length) and visible morphological features such as digit formation and eyelid morphogenesis. Additional features described here, including the acquisition of epidermal thickening, immunohistochemistry for Krt10, and expression of *Dkk4* as assessed by *in situ* hybridization, were used to refine stages 15 and 16 into substages. Stage 15b is distinguished from 15a by expression of *Dkk4*. (We note that stage 15a scRNAseq identifies a small number of cells that express *Dkk4*, but that are not detectable by *in situ* hybridization (Figs. 2b, 2c)). In the skin, stage 16a is distinguished from 15b by the appearance of epidermal thickening (and expression of *Dkk4* in the thickened regions). Stage 16b is distinguished from 16a by the appearance of some Krt10-positive cells in thick regions.

At stages 16a and 16b, thick regions have 3-5 layers of monomorphous, basaloid cells, whereas thin regions have 1-2 cell layers. Several observations indicate that the thick and thin regions are fundamentally different from epidermal stratification that normally occurs later in development: (1) suprabasal keratinocytes in both regions express a keratin, Krt5, that is normally limited to basal keratinocytes [2]; (2) cell proliferation as indicated by expression of Ki-67 occurs in all epidermal layers rather than being limited to basal keratinocytes; and (3) the keratin, Krt10, which is normally distributed uniformly in suprabasal keratinocytes at stage 18 and beyond (Fig. 1c), exhibits an unusual pattern of expression in thick regions towards the end of stage 16 (Figs. 1b and 1c, Supplementary Table 1).

Stage 17 is distinguished from stage 16b by fetal size, eyelid closure, and the initiation of hair follicle development (Supplementary Table 1). In addition, between stages 15b and 17, the size of the *Dkk4*+ expression domain becomes smaller and is eventually restricted to the hair bud (Supplementary Fig. 2). Changes in epidermal morphology and expression of *Dkk4* in stages 15 and 16a are not observed at the corresponding stages in laboratory mice. Morphological differences between these previously unknown aspects of cat epidermal development at stages 15 and 16, and normal hair follicle development and epidermal stratification that occurs later in other mammals (including the cat) are summarized in Supplementary Table 2.

We assessed the effect of *Taqpep* genotype on epidermal topology by morphometric reconstruction from serial sections from the flank (Fig. 1d) and the dorsal neck (Supplementary Fig. 1) of $Ta^M$/- and $Ta^b/Ta^b$ embryos. The orientation of tabby stripes with respect to the body axis differs between the flank and the neck, but in both cases, the patterns of fetal epidermal topology and their response to variation in *Taqpep* genotype correspond to stereotypic postnatal color patterns. Specifically, in fetal skin from the flank, thick epidermal regions are organized into columns that are perpendicular to the body axis (black bars, Fig. 1d); in fetal skin from the dorsal neck, thick epidermal regions lie in narrow, vertical columns that are parallel to the body axis (black bars, Supplementary Fig. 1). In both cases, the thick regions are expanded in $Ta^b/Ta^b$ embryos.

<u>Supplementary Note 2: Single-cell gene expression experiments and analyses</u>

We constructed and analyzed a total of 9 scRNAseq libraries from fetal skin, of which 3 are described in detail here. Each of the 3 libraries consisted of >4400 cells, with median gene and unique transcript counts ranging from 2,570-4,119 and 9,419-18,244, respectively, and 24,164-25,514 genes detected (Supplementary Table 4).

For the library from stage 16a, we used *k*-means clustering and gene markers to assign the identity of specific cell clusters (Supplementary Table 3, Fig. 2a). At *k*=7, those clusters correspond to myoblasts, neural crest, dendritic cells, endothelium, dermal fibroblasts, and two types of keratinocytes, a predominant basal population distinguished by high levels of *Krt5* expression, and a smaller differentiated population expressing non-basal markers, including *Keratinocyte Differentiation Associated Protein*, *Keratin 7*, and *Beta-defensin 1*. At *k*=8, vascular endothelium can be distinguished from lymphatic endothelium. At *k*=9, the basal keratinocytes cluster into two subpopulations based on differential expression of 604 genes, and in which expression of *Dkk4* and *Wif1* are the most significant, p=6.50e-46 and p=5.81e-50, respectively (negative binomial exact test, Supplementary Table 5).

For stage 15b, a subset of basal keratiocytes had a distinct gene expression signature characterized by the expression of *Engrailed* (*En1*) and *HoxC* genes, likely to reflect cells from the developing limb bud. The *En1*/*HoxC* signature was not observed at stages 15a or 16a, and therefore this population of cells was excluded from the stage15b differential expression analysis.

For stage 15a, *k*-means clustering (*k*=10) did not resolve subpopulations of basal keratinocytes, likely due to the relatively small number of *Dkk4*-positive cells (Figs. 2b, 2c). To recognize and distinguish *Dkk4*-positive and *Dkk4*-negative basal keratinocytes at stages 15a and 15b, we used an empiric threshold for *Dkk4* expression, e.g. all cells in the data set demonstrating 4-fold elevated *Dkk4* expression (relative to all cells, and equivalent to > 2 *Dkk4* UMI counts per cell) were categorized as the *Dkk4*-positive population, and all basal keratinocytes (*Krt5*-positive; *Krtdap*-negative) with <4-fold *Dkk4* levels (relative to all cells) were categorized as the *Dkk4*-negative population. We tested thresholds of 2-, 4-, 8-, and 16-fold, and observed little impact on the number of basal keratinocytes deemed *Dkk4*-positive and *Dkk4*-negative at each of the 3 stages (Supplementary Table 8). The UMAP projections for stages 15a and 15b (Fig. 2b) are based on aggregation of the stage 15a and 15b expression matrices.

Supplementary Table 5 displays expression levels, fold-change between *Dkk4*-positive and *Dkk4*-negative cells, and false discovery rates (FDR) for all genes. For comparison of genes across the different stages, we used a differential expression threshold rather than an FDR threshold, since FDR is biased by the population size differences at each time point. (At stage 15a, comparison of 59 *Dkk4*-positive to 1569 *Dkk4*-negative cells yields 13 of 21761 genes with an FDR < 0.05, but a 2-fold differential expression threshold yields 928 genes, of which 184 are in common with stages 15b and 16a (Fig. 3a, 3b)). The stage-specific gene lists used for Figure 3 are based on (1) a >2-fold expression difference between basal keratinocyte subpopulations; and (2) a median normalized average of >1 transcript per 10 cells in at least one basal keratinocyte population.

Supplementary Note 3: Genetic evaluation of *Ticked*

Our initial evaluation of *Dkk4* as a candidate gene for *Ticked* was based on its genetic map location [3,4] and a survey of existing genome sequence data from the 99 Lives collection [5,6] that identified the p.Ala18Val and p.Cys63Tyr variants. We ascertained additional DNA samples from cat shows and breeders, and carried out association, haplotype, and segregation analysis to further evaluate the *Dkk4* variants.

For some breeds, the breed standard is such that *Ticked* is either required (Abyssinian, Singapura) or under strong selection (Burmese). In the Abyssinian and Singapura breeds, a distinctive characteristic is prominent banding within individual hairs, uniformly distributed along most of the body surface. Among 37 Abyssinian and 26 Singapura cats, all carried the p.Ala18Val or p.Cys63Val variants, the vast majority as homozygotes or compound heterozygotes (Table 1, Supplementary Table 7). In the Burmese breed, distinctive characteristics are point coloration, caused by a temperature sensitive *Tyrosinase* variant [7], and an otherwise uniform appearance without tabby pattern or hair banding, caused by a loss-of-function *Agouti* allele [8]. However, in cats with a dark base color, “ghost tabby markings” are sometimes visible; consequently, *Ticked* is selected by Burmese breeders, not for its ability to promote hair banding, but for its epistasis over *Tabby* [3,9]. Among 13 Burmese cats, 11 were homozygous for the *Dkk4* p.Ala18Val variant, 1 was heterozygous, and 1 was homozygous for the ancestral allele (Table 1, Supplementary Table 7).

For other breeds including the Egyptian Mau, Ocicat, and Bengal cat, tabby markings are required, therefore *Ticked* should not be present due to its epistasis over *Tabby* [9]. In 31 such animals, none carried a derivative *Dkk4* allele (Table 1, Supplementary Table 7). Finally, in feral cats and in some breeds such as the Oriental Longhair (OLH) and Oriental Shorthair (OSH), both Ticked and non-Ticked phenotypes are observed. In this group, we found that 29 of 29 Ticked cats were heterozygous or homozygous for the *Dkk4* p.Ala18Val variant, while no *Dkk4* deleterious variants were observed in 209 non-Ticked cats (Table 1, Supplementary Table 7).

We used genome sequence data from 57 cats in the 99 Lives dataset [6] to evaluate shared haplotypes surrounding the p.Ala18Val (n=12) and p.Cys63Tyr (n=6) variants (Fig. 4b, Supplementary 4a). As with the linkage interval, both haplotypes are long, 6.08 and 8.95 Mb, respectively, likely due to suppressed recombination near the chrB1 centromere. The haplotype carrying the p.Cys63Tyr variant was not observed in any cats without the variant, and reflects the results from the association analysis (Supplementary Table 7), in which p.Cys63Tyr is restricted to the Abyssinian and Singapura breeds. By contrast, the p.Ala18Val variant is more broadly distributed in Abyssinian, Singapura, Burmese, OSH, and non-breed cats (Supplementary Table 7), and the haplotype upon which the p.Ala18Val variant exists was observed several times in cats that did not carry the variant (Supplementary Fig. 4a, Birman*).

We evaluated Mendelian transmission and allelic interactions in an OSH pedigree (Supplementary Fig. 4b), and observed that the p.Ala18Val variant cosegregated perfectly with the Ticked phenotype (13 Ticked, 8 non-Ticked). In a Singapura pedigree that segregated both alleles (Supplementary Fig. 4c, the Ticked phenotype was observed in p.Ala18Val homozygotes (n=6), p.Cys63Tyr homozygotes (n=1), and compound heterozygotes (n=7).

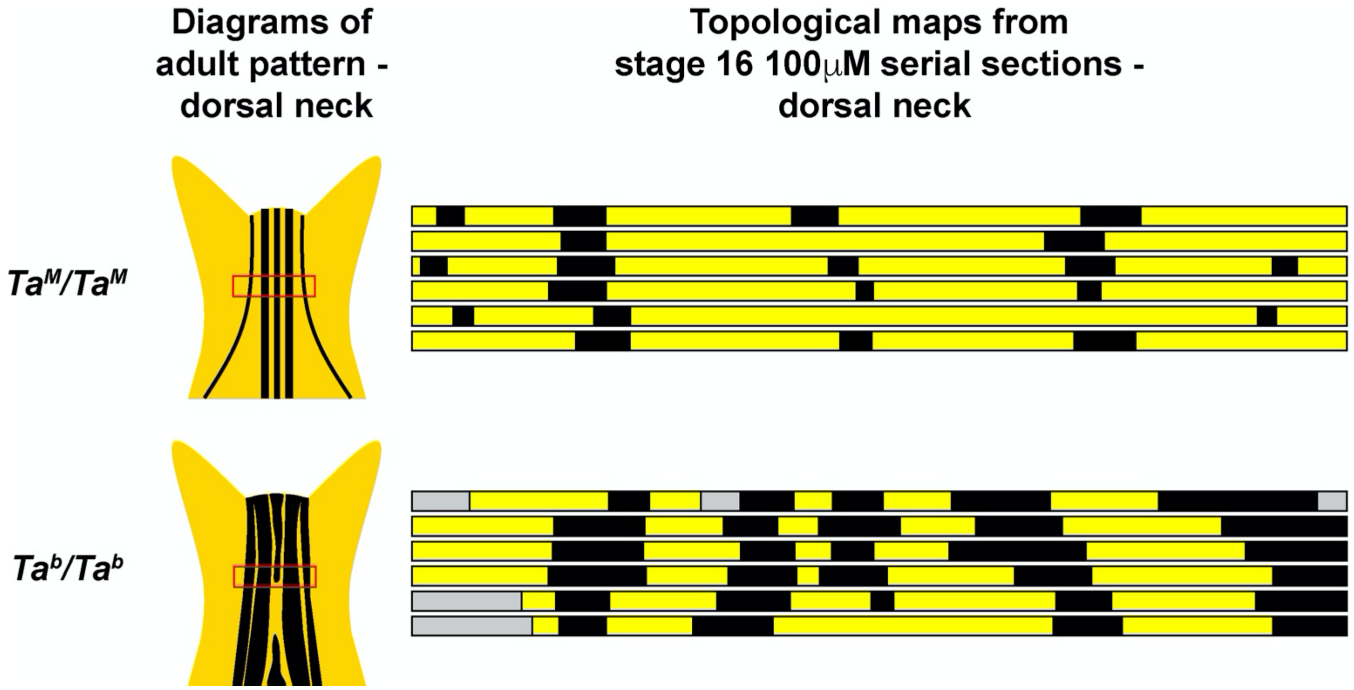


**Supplementary Fig. 1. Topological maps of dorsal neck skin from Stage 16.** *$Ta^M/Ta^M$* and *$Ta^b/Ta^b$* embryos (thin epidermis, yellow; thick epidermis, black; no data, grey). Cartoons show adult pigmentation pattern on the dorsal neck (red box indicates anatomic location of map).

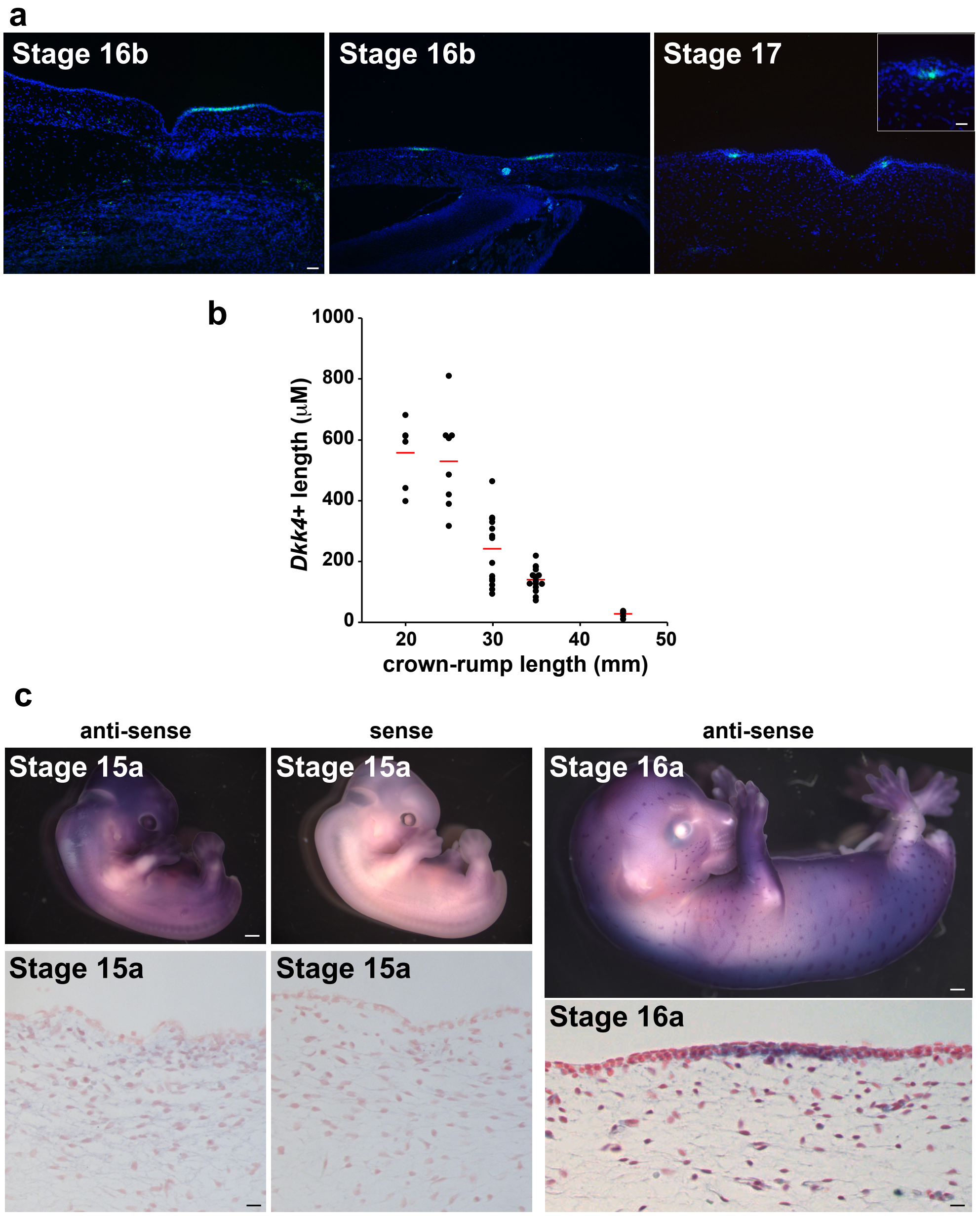


**Supplementary Fig. 2. *Dkk4* expression in embryonic cat skin.** (**a**) *Dkk4* expression (green) in sections of embryonic cat skin (DAPI, blue). 1-3 *Dkk4*-positive cells are found in sections of developing hair buds at stage 17 (right panel, inset). (**b**) Length of *Dkk4*-positive regions from sections of embryonic skin at different developmental time points (crown-rump length is a surrogate for stage of embryonic development, Supplementary Table 1; length of 6-16 *Dkk4*-positive regions measured from at least three sections at each developmental time point, red bar denotes mean). (**c**) *Dkk4* expression (purple) in stage 15a and stage 16a embryos (top panels; sense control probe does not stain embryo). Sections of skin (bottom panels) from embryos shown in top panels are stained with nuclear fast red. Scale bars: a, 50 μM; b inset, 25 μM; c top panels, 1mm; c bottom panels 25 μM.

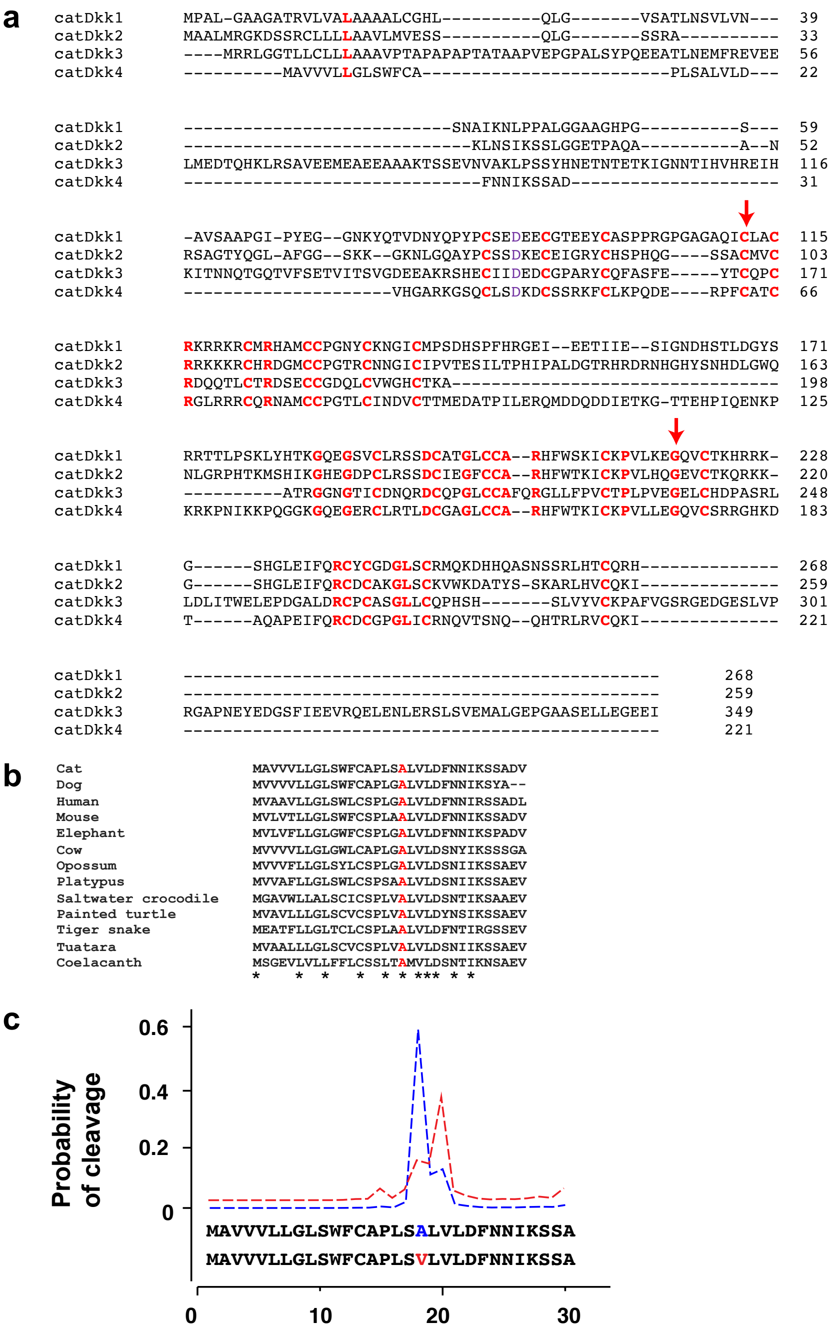


**Supplementary Fig. 3. Evolutionary constraint and functional prediction for cat Dkk4 variants associated with *Ticked*.** (**a**) A protein alignment of cat Dkk paralogs, with conserved residues colored red, and arrows highlighting the position of one of the cat *Dkk4* mutations (p.Cys63Tyr) and the position of the potentially deleterious residue in tiger (p.Gly173Glu). **(b)** Vertebrate Dkk4 alignment with the residue that harbors the other cat *Dkk4* mutation (p.Ala18Val) colored red. **(c)** Signal peptide cleavage site efficiencies predicted by SignalP5.0 for p.Ala18 (blue) and p.Val18 (red) variants.

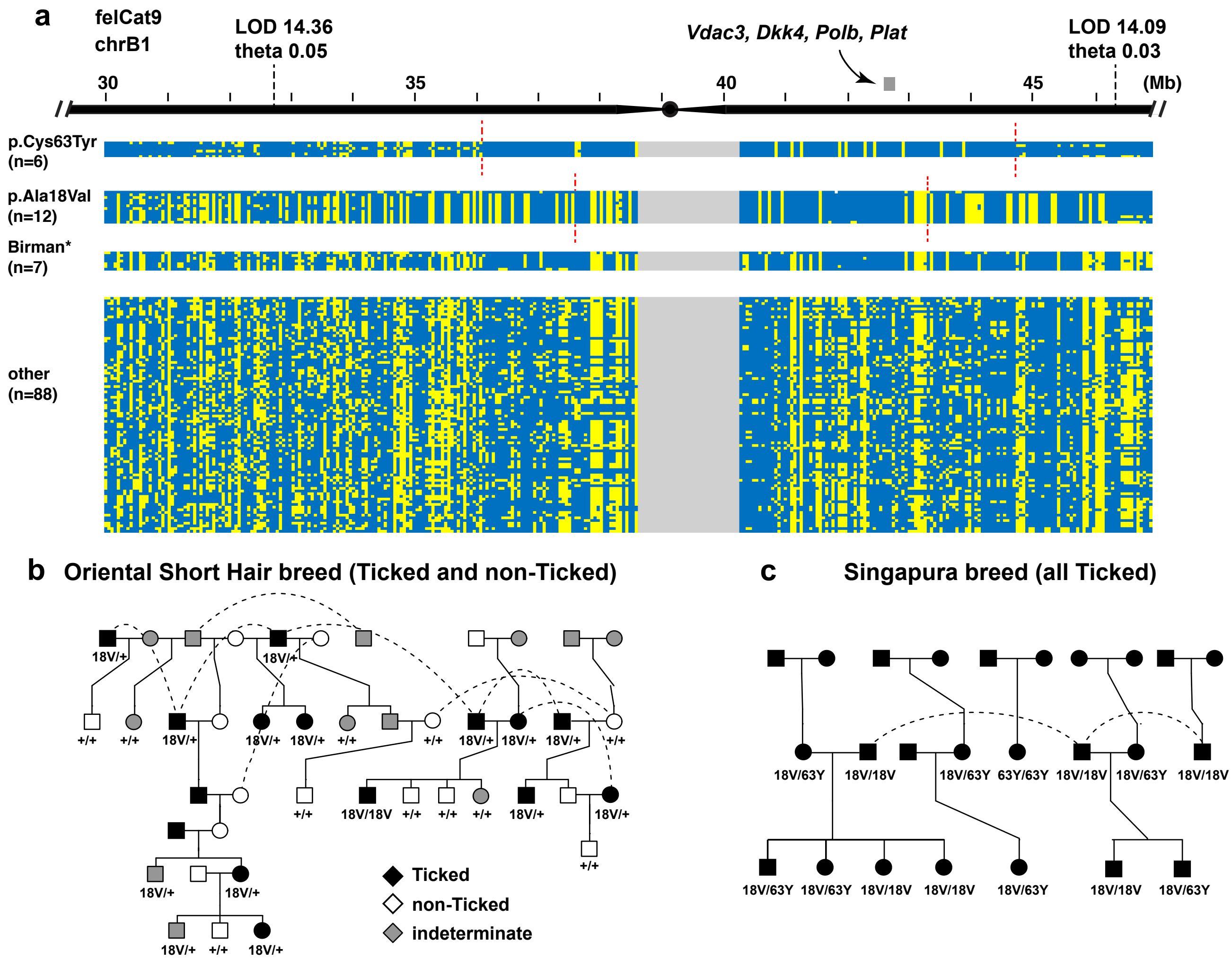


**Supplementary Fig. 4. *Dkk4* variants occur on extended haplotypes and co-segregate with *Ticked* in pedigrees.** (**a**) Haplotypes (rows) were inferred using Beagle (v4.1) for 57 cats in the 99 Lives data set, with SNPs (columns) indicated as reference (blue) or alternate (yellow) alleles. Broken black and red lines delineate the *Ticked* linkage interval and inferred ancestral recombination breakpoints on *Ticked* haplotypes, respectively. The grey bar indicates a region containing four genes that are differentially expressed between stage 16a basal keratinocyte subpopulations. The Birman* haplotype group consists of Birman (n=5), Devon Rex (n=1), and Domestic Short Hair (n=1) haplotypes, which are similar to the shared p.Ala18Val haplotype but do not carry the p.Ala18Val allele. **(b)** An Oriental Shorthair pedigree, demonstrating cosegregation of the Ticked pattern and the p.Ala18Val allele. **(c)** A Singapura pedigree, demonstrating fixation of the Ticked pattern as a consequence of p.Ala18Val and p.Cys63Tyr segregation.

```
Cheetah               MAVVFLLGLSWFCAPLSALVLDFNNIKSSANVHRAQKGSQCLSDKDCSSRKFCLKPQDER    60
Cougar                MAVVFLLGLSWFCAPLSALVLDFNNIKSSADVHGAQKGSQCLSDKDCSSRKFCLKPQDER    60
Tiger                 MAVVVLLGLSWLCAPLSALVLDFNNIKSSADVQGARKGSQCLSDKDCSSRKFCLKPQDER    60
Fishing cat           MAVVVLLGLSWFCAPLSALVLDFNNIKSSADVHGARKGSQCLSDKDCSSRKFCLKPQEER    60
Jaguar                MAVVVLLGLSWFCAPLSALVLDFNNIKSSADVHGARKGSQCLSDKDCSSRKFCLKPQDER    60
Lion                  MAVVVLLGLSWFCAPLSALVLDFNNIKSSADVHGARKGSQCLSDKDCSSRKFCLKPQDER    60
Snow leopard          MAVVVLLGLSWFCAPLSALVLDFNNIKSSADVHGARKGSQCLSDKDCSSRKFCLKPQDER    60
Leopard               MAVVVLLGLSWFCAPLSALVLDFNNIKSSADVHGARKGSQCLSDKDCSSRKFCLKPQDER    60
Asian leopard cat     MAVVVLLGLSWFCAPLSALVLDFNNIKSSADVHGARKGSQCLSDKDCSSRKFCLKPQDER    60
Clouded leopard       MAVVVLLGLSWFCAPLSALVLDFNNIKSSADVHGARKGSQCLSDKDCSSRKFCLKPQDER    60
Pallas cat            MAVVVLLGLSWFCAPLSALVLDFNNIKSSADVHGARKGSQCLSDKDCSSRKFCLKPQDER    60
Spanish lynx          MAVVVLLGLSWFCAPLSALVLDFNNIKSSADVHGARKGSQCLSDKDCSSRKFCLKPQDER    60
Pampas cat            MAVVVLLGLSWFCAPLSALVLDFNNIKSSADVHGARKGSQCLSDKDCSSRKFCLKPQDER    60
Canadian lynx         MAVVVLLGLSWFCAPLSALVLDFNNIKSSADVHGARKGSQCLSDKDCSSRKFCLKPQDER    60
Domestic cat          MAVVVLLGLSWFCAPLSALVLDFNNIKSSADVHGARKGSQCLSDKDCSSRKFCLKPQDER    60
Sand cat              MAVVVLLGLSWFCAPLSALVLDFNNIKSSADVHGARKGSQCLSDKDCSSRKFCLKPQDER    60
Jungle cat            MAVVVLLGLSWFCAPLSALVLDFNNIKSSADVHGARKGSQCLSDKDCSSRKFCLKPQDER    60
Black-footed cat      MAVVVLLGLSWFCAPLSALVLDFNNIKSSADVHGARKGSQCLSDKDCSSRKFCLKPQDER    60
                      ****.******:******************:*: *:*********************:**

Cheetah               PLCATCRGLRRRCQRNAMCCPGILCMNDVCTAMEDATPILERQMDDQDDIETKGTTEHPI    120
pumaDkk4              PFCATCRGLRRRCQRNAMCCPGTLCMNDVCTAMEDATPILERQMDDQDDIETKGTTEHPI    120
tigerDkk4             PFCATCRGLRRRCQRNAMCCPGTLCMNDVCTTMEDATPILERQMDDQDDIETKGTTEHPI    120
fishing_catDkk4       PFCATCRGLRRRCQRNAMCCPGTLCMNDVCTTMEDATPILERQMDDQDDIETKGTTEHPI    120
jaguarDkk4            PFCATCRGLQRRCQRNAMCCPGTLCMNDVCTTMEDATPILERQMDDQDDIETKGTTEHPI    120
lionDkk4              PFCATCRGLRRRCQRNAMCCPGTLCMNDVCTTMEDATPILERQMDDQDDIETKGTTEHPI    120
snow_leopardDkk4      PFCATCRGLRRRCQRNAMCCPGTLCMNDVCTTMEDATPILERQMDDQDDIETKGTTEHPI    120
leopardDkk4           PFCATCRGLRRRCQRNAMCCPGTLCMNDVCTTMEDATPILERQMDDQDDIETKGTTEHPI    120
alcDkk4               PFCATCRGLRRRCQRNAMCCPGTLCMNDVCTTMEDATPILERQMDDQDDIETKGTTEHPI    120
cloudedDkk4           PFCATCRGLRRRCQRNAMCCPGTLCMNDVCTTMEDATPILERQMDDQDDIETKGTTEHPI    120
pallas_catDkk4        PFCATCRGLRRRCQRNAMCCPGTLCMNDVCTTMEDATPILERQMDDQDDIETKGTTEHPI    120
SpanishLynxDkk4       PFCATCRGLRRRCQRNAMCCPGTLCMNDVCTTMEDATPILERQMDDQDDIETKGTTEHPI    120
pampasDkk4            PFCATCRGLRRRCQRNAMCCPGTLCMNDVCTTMEDATPILERQMDDQDDIETKGTTEHPI    120
CanadianLynxDkk4      PFCATCRGLRRRCQRNAMCCPGTLCMNDVCTTMEDATPILERQMDDQDDIETKGTTEHPI    120
catDkk4               PFCATCRGLRRRCQRNAMCCPGTLCINDVCTTMEDATPILERQMDDQDDIETKGTTEHPI    120
sand_catDkk4          PFCATCRGLRRRCQRNAMCCPGTLCINDVCTTMEDATPILERQMDDQDDIETKGTTEHPI    120
chausDkk4             PFCATCRGLRRRCQRNAMCCPGTLCINDVCTTMEDATPILERQMDDQDDIETKGTTEHPI    120
nigripesDkk4          PFCATCRGLRRRCQRNAMCCPGTLCINDVCTTMEDATPILERQMDDQDDIETKGTTEHPI    120
                      *:*******:************ **:*****:****************************

cheetahDkk4           QENKPKRKPNIKKPQDGKGQEGERCLRTLDCGAGLCCARHFWTKICKPVLLEGQVCSRRG    180
pumaDkk4              QENKPKRKPNIKKPQDGKGQEGERCLRTLDCGAGLCCARHFWTKICKPVLLEGQVCSRRG    180
tigerDkk4             QENKPKRKPNIKKPQGGKGQEGERCLRTLDCGAGLCCARHFWTKICKPVLLEEQVCSRRG    180
fishing_catDkk4       QENKPKRKPNIKKPQGGKGQEGERCLRTLDCGAGLCCARHFWTKICKPVLLEGQVCSRRG    180
jaguarDkk4            QENKPKRKPNIKKPQGGKGQEGERCLRTLDCGAGLCCARHFWTKICKPVLLEGQVCSRRG    180
lionDkk4              QENKPKRKPNIKKPQGGKGQEGERCLRTLDCGAGLCCARHFWTKICKPVLLEGQVCSRRG    180
snow_leopardDkk4      QENKPKRKPNIKKPQGGKGQEGERCLRTLDCGAGLCCARHFWTKICKPVLLEGQVCSRRG    180
leopardDkk4           QENKPKRKPNIKKPQGGKGQEGERCLRTLDCGAGLCCARHFWTKICKPVLLEGQVCSRRG    180
alcDkk4               QENKPKRKPNIKKPQGGKGQEGERCLRTLDCGAGLCCARHFWTKICKPVLLEGQVCSRRG    180
cloudedDkk4           QENKPKRKPNIKKPQGGKGQEGERCLRTLDCGAGLCCARHFWTKICKPVLLEGQVCSRRG    180
pallas_catDkk4        QENKPKRKPNIKKPQGGKGQEGERCLRTLDCGAGLCCARHFWTKICKPVLLEGQVCSRRG    180
SpanishLynxDkk4       QENKPKRKPNIKKPQGGKGQEGERCLRTLDCGAGLCCARHFWTKICKPVLLEGQVCSRRG    180
pampasDkk4            QENKPKRKPNIKKPQGGKGQEGERCLRTLDCGAGLCCARHFWTKICKPVLLEGQVCSRRG    180
CanadianLynxDkk4      QENKPKRKPNIKKPQGGKGQEGERCLRTLDCGAGLCCARHFWTKICKPVLLEGQVCSRRG    180
catDkk4               QENKPKRKPNIKKPQGGKGQEGERCLRTLDCGAGLCCARHFWTKICKPVLLEGQVCSRRG    180
sand_catDkk4          QENKPKRKPNIKKPQGGKGQEGERCLRTLDCGAGLCCARHFWTKICKPVLLEGQVCSRRG    180
chausDkk4             QENKPKRKPNIKKPQGGKGQEGERCLRTLDCGAGLCCARHFWTKICKPVLLEGQVCSRRG    180
nigripesDkk4          QENKPKRKPNIKKPQGGKGQEGERCLRTLDCGAGLCCARHFWTKICKPVLLEGQVCSRRG    180
                      ***************.************************************ *******

cheetahDkk4           HKDTAQAPEIFQRCDCGPGLICRNQVTGNQQHTRLRVCQKI--  221
pumaDkk4              HKDTAQAPEIFQRCDCGPGLICRNQVTSNQQHTRLRVCQKI--  221
tigerDkk4             HKDTAQAPEIFQRCDCGPGLICRNQVTGNQQHTRLRVCQKI--  221
fishing_catDkk4       HKDTAQAPEIFQRCDCGPGLICRNQVTGNQQHTRLRVCQKI--  221
jaguarDkk4            HKDTAQAPEIFQRCDCGPGLICRNQVTGNQQHTRLRVCQKI--  221
lionDkk4              HKDTAQAPEIFQRCDCGPGLICRNQVTGNQQHTRLRVCQKNLN  223
snow_leopardDkk4      HKDTAQAPEIFQRCDCGPGLICRNQVTGNQQHTRLRVCQKI--  221
leopardDkk4           HKDTAQAPEIFQRCDCGPGLICRNQVTGNQQHTRLRVCQKI--  221
alcDkk4               HKDTAQAPEIFQRCDCGPGLICRNQVTGNQQHTRLRVCQKI--  221
cloudedDkk4           HKDTAQAPEIFQRCDCGPGLICRNQVTGNQQHTRLRVCQKI--  221
pallas_catDkk4        HKDTAQAPEIFQRCDCGPGLICRNQVTGNQQHTRLRVCQKI--  221
SpanishLynxDkk4       HKDTAQAPEIFQRCDCGPGLICRNQVTGNQQHTRLRVCQKI--  221
pampasDkk4            HKDTAQAPEIFQRCDCGPGLICRNQVTGNQQHTRLRVCQKI--  221
CanadianLynxDkk4      HKDTAQAPEIFQRCDCGPGLICRNQVTSNQQHTRLRVCQKI--  221
catDkk4               HKDTAQAPEIFQRCDCGPGLICRNQVTSNQQHTRLRVCQKI--  221
sand_catDkk4          HKDTAQAPEIFQRCDCGPGLICRNQVTSNQQHTRLRVCQKI--  221
chausDkk4             HKDTAQAPEIFQRCDCGPGLICRNQVTGNQQHTRLRVCQKI--  221
nigripesDkk4          HKDTAQAPEIFQRCDCGPGLICRNQVTGNQQHTRLRVCQKI--  221
                      ***************************.************
```

**Supplementary Fig. 5. Protein alignment of predicted Dkk4 protein sequence from 29 felid species.** A summary of derived variants and predicted deleteriousness is given in Supplementary Table S9.

**Supplementary Table 1. Stages of embryonic and fetal development in the domestic cat.**

| Prenatal stage (cat)[a, b] | Days post-coitus[a] | Crown-rump length (mm)[b] | Prenatal stage (mouse)[c] | Anatomic features (cat)[a,b] | Histologic features (cat)[b] | Molecular features (cat)[b] |
|---|---|---|---|---|---|---|
| 13 | 19-21 | 8-11 | E11 | | monolayer of epidermal cells | |
| 14 | 21-23 | 11-15 | E12 | | | |
| 15a | 23-25[a,b] | 15-17 | E14 | finger web resorption | | |
| 15b | 23-25[a,b] | 18-21 | E14 | toe web resorption | | *Dkk4*-positive domains in epidermis |
| 16a | 25-28[a,b] | 22-26 | E15 | eye lid closure begins | “thick” epidermal domains | *Dkk4*-positive “thick” domains |
| 16b | 25-28[a,b] | 27-37 | E15 | | | Krt10-positive cells in “thick” domains |
| 17 | 28-32 | 38-47 | E16 | eyelid closure complete | hair follicle placodes and buds | *Dkk4*-positive cells in hair buds |
| 18 | 32-38 | 48-60 | | | epidermis uniform thickness | Krt10-positive uniformly expressed in epidermis |
| 19 | 38-44 | 61-80 | E17 | | | |
| 20 | 44-48 | 81-90 | | | | |
| 21 | 48-60 | 91-109 | | hair eruption; pigment pattern visible | pigmented hair bulbs | |
| 22 | 60-66 | 110-130 | P4-6 | | | |

[a] Based on Knospe et al. [1]
[b] Based on data reported here
[c] Based on Kaufman et al. [10]

**Supplementary Table 2. Histologic and molecular features of thick epidermal domains in cat as compared to developing hair follicle placodes and "normal" epidermal stratification.**

| | "Thick" epidermal domain (cat[a]) | Hair follicle placode (mouse, cat[a]) | Developing epidermis (mouse, cat[a]) |
|---|---|---|---|
| **Tissue organization** | 3-5 layers of uniformly spaced, basaloid cells | Single layer of crowded, basal cells below second layer of basal-like cells, intermediate cell layer and periderm [11]* | Single layer of uniformly spaced basal cells below intermediate cell layer and periderm [12]; intermediate cell layer differentiates into spinous, granular, cornified layers; periderm sheds [12] |
| **Cellular morphology** | Round, basaloid cells uniformly spaced | Vertically polarized, basal cells undergo compaction [11,13]; round, basaloid and intermediate cells | Round, basal and intermediate cells uniformly spaced [12]; differentiated epidermal cells flatten |
| **Domain width** | 50-500 μM (~20-200 cells); $Ta^b/Ta^b > Ta^M/-$ | ~50 μM (~5-20 cells) [11] | |
| **Domain depth** | No invagination into dermis | Invaginates into dermis [11]* | |
| **Proliferation** | Basal and supra-basal cells proliferate (31% Ki67-positive) | Small fraction proliferating cells (~8%, thickening due to cell migration) [13]; basal cells proliferate; supra-basal cells proliferate during invagination [13,14] | Basal and few supra-basal cells located just above the basal cell layer proliferate [12,15] |
| **Keratin expression pattern** | Krt5-positive basal and supra-basal cells; Krt10-positive cells lie above 3-5 layers of Krt5-positive cells | Krt5-positive basal and basaloid cells; Krt10-positive intermediate cell layer; Krt8/18-positive periderm [12] | Krt5-positive basal cells [12]; Krt1/10-positive supra-basal cells [12,15]; Krt8/18-positive periderm [12] |
| **Dermal cell arrangement** | Dermal cells evenly distributed | Dermal cells cluster below placode [11] | Dermal cells evenly distributed |

[a]Data in cat from this study.

**Supplementary Table 3. Gene markers used for identification of UMAP clusters.**

| Cell population | Key genes | Reference |
|---|---|---|
| Macrophage/dendritic cells | *Cd163*, *C1qa*, *Cd14* | [16,17] |
| Neural Crest | *Fabp7*, *Sox10*, *Foxd3* | [18] |
| Stratified epithelium/periderm | *Krtdap*, *Rhov*, *Defb1* | [19] |
| Basal keratinocytes | *Krt5*, *Tp63*, *Kremen2* | [20] |
| Dermal fibroblasts | *Lum*, *Twist1*, *Twist2* | [20] |
| Myoblasts | *Myf5*, *Myod1*, *Msc* | [21,22] |
| Endothelium | *Pecam1*, *Cdh5*, *Tie1* | [23] |
| Vascular endothelium | *Cd34*, *Esam*, *Fam198b* | [24] |
| Lymphatic endothelium | Ccl21, Mmrn1, Igf1 | [25] |

**Supplementary Table 4. scRNAseq summary statistics.**

| | Embryonic stage | | |
|---|---|---|---|
| | 15a | 15b | 16a |
| ID | C16 | C14 | C64 |
| 10x Genomics Chemistry | v3 | v3 | v2 |
| Total reads | 455,308,860 | 450,083,295 | 404,331,199 |
| % mapped to genome | 87.9 | 87.3 | 87.9 |
| % mapped to transcriptome | 58.9 | 60.1 | 63.5 |
| Estimated cell count | 5,493 | 4,454 | 4,617 |
| Mean reads per cell | 82,889 | 101,051 | 87,574 |
| Median genes per cell | 4,119 | 4,180 | 2,570 |
| Median UMI per cell | 18,244 | 17,966 | 9,419 |
| Total genes detected | 25,514 | 25,222 | 24,164 |
| Estimated cell number | 5,493 | 4,454 | 4,617 |
| Keratinocyte cell number | 1,791 | 2,425 | 2,515 |
| Keratinocyte cell fraction | 0.33 | 0.54 | 0.54 |

**Supplementary Table 5. List of differentially expressed genes from scRNAseq and overlap with hair follicle placode. (separate Excel spreadsheet)**

### Supplementary Table 6. *Dkk4* coding variants detected in 57 cats[a].

| felCat9 coordinate | cDNA variant[b] | Amino acid variant[b] | Allele freq (n=57)[c] | Abyssinian allele freq (n=4)[c] | CADD phred |
|---|---|---|---|---|---|
| chrB1:42620835 | c41t | A18V | 0.10 | 0.13 | 22.9 |
| chrB1:42621481 | g188a | C63Y | 0.05 | 0.75 | 21.4 |
| chrB1:42621505 | g212a | R71K | 0.14 | 0.75 | 5.8 |
| chrB1:42621551 | a258g | I86M | 0.01 | 0 | 0.3[d] |
| chrB1:42622162 | a395g | K132R | 0.12 | 0 | 14.2 |
| chrB1:42623424 | t602c | I201T | 0.66 | 0.13 | 16.0[e] |
| chrB1:42623444 | a622g | S208G | 0.01 | 0 | 0.2 |
| chrB1:42623455 | a633c | Q211H | 0.01 | 0 | 0.2[d] |

[a] All variants predicted to alter amino acids in Dkk4 are shown. In 57 cat genome sequences from the 99 Lives collection, there are an additional 3 variants in *Dkk4* that are synonymous, and with CADD scores of *Q*<10. No variants predicted to alter amino acids were identified in Abyssinians for the 3 other genes that are differentially expressed and overlap with the *Ticked* linkage interval, *Plat*, *Polb*, and *Vdac3*.

[b] Ancestral allele is shown as the reference allele.

[c] Allele frequencies for the derived allele.

[d] Reference base between cat and human differs.

[e] Adjacent nucleotide position differs between cat and human, affecting the reference alternate codon, making CADD score interpretation unreliable.

**Supplementary Table 7.** ***Dkk4*** **alleles categorized according to breed and phenotype[a].**

| | *Dkk4* genotype | | | | | | |
|---|---|---|---|---|---|---|---|
| Breed and phenotype | +/+ | +/18V | +/63Y | 18V/18V | 18V/63Y | 63Y/63Y | Total |
| Abyssinian (Ticked) | 0 | 0 | 1 | 0 | 2 | 34 | 37 |
| Singapura (Ticked) | 0 | 1 | 1 | 4 | 16 | 4 | 26 |
| Burmese (Ticked) | 1 | 1 | 0 | 11 | 0 | 0 | 13 |
| Mau (non-Ticked) | 8 | 0 | 0 | 0 | 0 | 0 | 8 |
| Ocicat (non-Ticked) | 13 | 0 | 0 | 0 | 0 | 0 | 13 |
| Bengal (non-Ticked) | 10[b] | 0 | 0 | 0 | 0 | 0 | 10[b] |
| OSH and OLH Ticked | 0 | 23 | 0 | 2 | 0 | 0 | 25 |
| OSH and OLH non-Ticked | 21 | 0 | 0 | 0 | 0 | 0 | 21 |
| Non-breed cats Ticked | 0 | 4 | 0 | 0 | 0 | 0 | 4 |
| Non-breed cats non-Ticked | 188 | 0 | 0 | 0 | 0 | 0 | 190 |

[a] The p.Ala18Val and p.Cys63Tyr variants are indicated as 18V and 63Y, respectively. The table includes data from 18 breed cats in the 99Lives dataset (4 Abyssinian, 3 Egyptian Mau, 1 Ocicat, 5 Bengal, 5 Burmese), for which photographs were not available, and for which phenotype was inferred based on breed identity; all other data is based on samples that we ascertained and collected. Of the Oriental Shorthair (OSH) and Oriental Longhair (OLH), 15 had an indeterminate phenotype for which we could not infer *Ticked* genotype, and one was excluded due to an inconsistency in Mendelian transmission.

[b] Data shown are from targeted genotyping or high-coverage WGS. In additional data from low-coverage (0.1x – 0.5x) WGS on 526 Bengal cats, neither *Dkk4* variant was observed.

**Supplementary Table 8. Basal keratinocyte subpopulation cell number at different *Dkk4* expression thresholds.**

| Stage | Subpopulation | Expression Threshold | | | |
|---|---|---|---|---|---|
| | | 2-fold | 4-fold | 8-fold | 16-fold |
| 15a | *Dkk4*- | 1641 | 1569 | 1670 | 1682 |
| 15a | *Dkk4*+ | 77 | 59 | 48 | 36 |
| 15b | *Dkk4*- | 824 | 865 | 916 | 980 |
| 15b | *Dkk4*+ | 758 | 717 | 666 | 602 |
| 16a | *Dkk4*- | 846 | 931 | 1018 | 1193 |
| 16a | *Dkk4*+ | 1216 | 1131 | 1044 | 869 |

**Supplementary Table 9. *Dkk4* variants and predicted impact (CADD) in the Felidae.**

| Position | Ref | Alt | Amino acid | Species | CADD |
|---|---|---|---|---|---|
| 42377041 | A | G | p.A2V | Tigrina | 3.393 |
| 42377033 | C | A | p.V5F | Puma, Jaguarundi, Cheetah | 0.736 |
| 42376989 | C | G | p.D58E | Fishing cat | 14.5 |
| 42377012 | G | A | p.F12L | Tiger | 10.88 |
| 42376955 | C | T | p.D31N | Cheetah | 14.02 |
| 42376947 | A | G | p.H32Q | Tiger | 0.68 |
| 42376946 | C | T | p.G33R | Cheetah | 10.84 |
| 42376939 | C | T | p.R35Q | Puma, Jaguarundi, Cheetah | 9.835 |
| 42375756 | G | T | p.F62L | Cheetah | 11.17 |
| 42375733 | C | T | p.R70Q | Jaguar | 1.566 |
| 42375694 | G | A | p.T83I | Cheetah | 8.476 |
| 42374918 | G | T | p.M86I | Domestic cat, Sand cat, Black-Footed cat, Jungle cat | 4.527 |
| 42374902 | T | C | p.T92A | Puma, Jaguarundi, Cheetah, Bay cat, Asian gold., Marbled cat | 0.571 |
| 42374889 | G | C | p.A96G | Jaguarundi | 4.66 |
| 42374887 | T | C | p.T97A | Serval | 0.771 |
| 42374845 | G | T | p.E111K | Flat-headed cat | 0.108 |
| 42374769 | C | T | p.G136D | Puma, Jaguarundi, Cheetah | 3.221 |
| 42374318 | G | C | p.A153S | Flat-headed cat | 10.67 |
| 42374257 | C | T | p.G173E | Tiger | 26.9 |
| 42374174 | G | T | p.I201V | Geoffroys cat,Tigrina | 2.552 |
| 42374153 | T | C | p.G208S | Domestic cat, Sand cat, Puma | 0.21 |
| 42374114 | T | C | p.I221V | Bay cat | 0.435 |
| 42374114 | T | TT | p.I221fsX4 | Lion | 19.07 |
